# The GT factor ZmGT-3b mediates growth–defense tradeoff by regulating photosynthesis and defense response^1^

**DOI:** 10.1101/2021.03.12.435151

**Authors:** Qianqian Zhang, E Lizhu, Weixing Dai, Mingliang Xu, Jianrong Ye

## Abstract

Plant growth and development face constant threat from various environmental stresses. Transcription factors (TFs) are crucial for maintaining balance between plant growth and defense. Trihelix TFs display multifaceted functions in plant growth, development, and responses to various biotic and abiotic stresses. Here, we explore the role of a trihelix TF, ZmGT-3b, in regulating the growth–defense tradeoff in maize (*Zea mays*). *ZmGT-3b* is primed for instant response to *Fusarium graminearum* challenge by implementing a rapid and significant reduction of its expression to suppress seedling growth and enhance disease resistance. *ZmGT-3b* knockdown led to diminished growth, but improved disease resistance and drought tolerance in maize seedlings. In *ZmGT-3b* knockdown seedlings, the chlorophyll content and net photosynthetic rate were strongly reduced, whereas the contents of major cell wall components, such as lignin, were synchronically increased. Correspondingly, *ZmGT-3b* knockdown specifically downregulated photosynthesis-related genes, especially *ZmHY5* (encoding a conserved central regulator of seedling development and light responses), but synchronically upregulated genes associated with secondary metabolite biosynthesis and defense-related functions. *ZmGT-3b* knockdown induced defense-related transcriptional reprogramming and increased biosynthesis of lignin without immune activation. These data suggest that ZmGT-3b is a regulator of plant growth–defense tradeoff that coordinates metabolism during growth-to-defense transitions by optimizing the temporal and spatial expression of photosynthesis- and defense-related genes.

**One-sentence summary:** ZmGT-3b regulates photosynthesis activity and synchronically suppresses defense response.

## Introduction

Members of the trihelix family, one of the first transcription factor (TF) families discovered in plants, are classified as GT factors. This is because the first core sequence isolated from these TFs was the GT element 5ʹ-GGTTAA-3ʹ. This element is sufficient for light induction of the light-induced genes, and trihelix family TFs bind specifically to GT elements (Green et al., 1987). The DNA-binding domain of these TFs features a typical trihelix structure (helix-loop-helix-loop-helix); that is, the helices form a bundle held together by a hydrophobic core that determines their specific binding to GT elements (Kaplan-Levy et al., 2012). The first trihelix TF discovered, GT-1 in pea (*Pisum sativum*), binds specifically to a GT element in the promoter of *rbcS-3A* and regulates its light-dependent expression (Green et al., 1987). To date, 30 and 31 trihelix TFs have been identified in Arabidopsis (*Arabidopsis thaliana*) and rice (*Oryza sativa*), respectively. Trihelix family members are grouped into five subfamilies, GT-1, GT-2, GT, SH4, and SIP, which were named after the first identified member of each subfamily. GT-1 proteins have one trihelix DNA-binding domain, GT-2 members have two DNA-binding domains, GT-1 and GT-2 members share high sequence similarity (Kaplan-Levy et al., 2012; Qin et al., 2014). *Arabidopsis* GT-1 directly activates the transcription of its target genes by stabilizing the TFIIA–TBP–TATA components of the pre-initiation complex (Le Gourrierec et al., 1999).

Initially, trihelix family members were found to participate in various plant developmental programs and light responses. Cloning and characterization of trihelix members from various plants has subsequently revealed their broad functional divergence in processes including the development of floral organs, embryos, seeds, stomata, and trichomes, the development of the seed scattering trait during crop domestication, and biotic and abiotic stress responses (Kaplan-Levy et al., 2012; Qin et al., 2014). The *Arabidopsis* GT-2 clade member PETAL LOSS (PTL) regulates floral organ morphogenesis (Brewer et al., 2004; Kaplan-Levy et al., 2014). *Arabidopsis* GT-2 Like 1 (AtGTL1) is a key regulator of ploidy-dependent trichome growth and drought tolerance; its mutants show increased polyploidy levels and reduced stomatal numbers (Breuer et al., 2009). Poplar (*Populus*) *PtaGTL1*, a homolog of *Arabidopsis AtGTL1*, is also involved in stoma and trichome development (Weng et al., 2012). The GT TFs ASIL1 and ASIL2 negatively regulate the expression of several genes encoding seed maturation proteins in *Arabidopsis* (Gao et al., 2009). The rice GT member SHA1 regulates the seed scattering process, a well-characterized domestication trait (Li et al., 2006). SHA1 promotes the function of the abscission layer in the pedicels of mature seeds (Lin et al., 2007). The biosynthesis of mixed-linkage glucan (MLG) depends on the biochemical activity of membrane-spanning cellulose synthase-like F/H (CSLF/H). The *Brachypodium distachyon* trihelix family TF BdTHX1 is involved in regulating MLG biosynthesis by controlling the transcription of *BdCSLF6* and the endotransglucosylase/hydrolase gene *BdXTH8* (Fan et al., 2018).

Trihelix TFs are also critical for plant responses to various biotic and abiotic stresses. The rice GT gene *OsRML1* is induced in response to infection by the fungal pathogen *Magnaporthe grisea* (Wang et al., 2004). Two *Arabidopsis* GT-1 clade members, *GT-3a* and *GT-3b*, function in the plant’s response to salt and pathogen stress (Park et al., 2004). AtGTL1 negatively regulates water use efficiency by modulating stomatal density; mutation of the encoding gene increases plant tolerance to drought stress (Yoo et al., 2010). Overexpression of the soybean (*Glycine max*) GT-2 genes *GmGT-2A* and *GmGT-2B* enhanced tolerance to salt, drought, and freezing stress in transgenic *Arabidopsis* (Xie et al., 2009). GTL1 is part of the MPK4-signaling cascade that coordinates pattern-triggered immunity (PTI) and effector-triggered immunity (ETI), as GTL1 positively regulates defense genes and inhibits factors that mediate plant growth and development; *gtl1* mutants are compromised in basal immunity, PTI, and ETI (Völz et al., 2018). However, ARABIDOPSIS SH4-RELATED3 (ASR3) is phosphorylated by MPK4 to negatively regulate flg22-induced gene expression and functions as a negative regulator of PTI (Li et al., 2015).

Plant growth and development are constantly affected by various environmental stresses. To survive under constantly changing environmental conditions, plants must maintain a dynamic growth–defense balance to allow optimal allocation of resources, which demands prioritization towards either growth or defense, depending on external and internal signals (Huot et al., 2014). The transcriptional regulation of gene expression is central to both plant development and responses to environmental stimuli. The induction of plant immune responses involves rapid transcriptional reprogramming that prioritizes defense-over growth-related cellular functions, which usually compromises vegetative tissue growth and yield (Alves et al., 2014). Transcriptional regulators are key components or master regulators of various signal transduction pathways that function during plant defense responses. TF activity alters the plant transcriptome, leading to molecular, metabolic, and phenotypic changes in favor of defense responses at the expense of normal growth (Alves et al., 2014). Various TFs are involved in regulating these processes by binding to specific *cis*-acting elements in the promoters of their target genes to activate or inhibit their transcription. The specific binding domain of each TF family binds to DNA *cis*-elements associated with responses to a specific environmental stress; these are key features distinguish one family from another (Riechmann et al., 2000; VanVerk et al., 2009; Mizoi et al., 2012). The best-known major TF families involved in plant defense responses are WRKYGQK (WRKY), basic leucine zipper (bZIP), myelocytomatosis (MYC), myeloblastosis (MYB), APETALA2/ ETHYLENE-RESPONSIVE ELEMENT BINDING FACTORS (AP2/EREBP), and NAM, ATAF and CUC (NAC) TFs (Riechmann et al., 2000; Alves et al., 2014).

The MYB TF family is one of the largest plant TF families. MYBs not only have multifaceted roles in plant growth and development, but also in many physiological and biochemical processes, especially the regulation of primary and secondary metabolism and responses to various biotic and abiotic stresses. MYBs may function in the crosstalk linking abiotic stress responses with lignin biosynthesis pathways (Baldoni et al., 2015). Lignin, a key secondary metabolite in plants, is a major structural component of the vascular cell wall, facilitates water transport, and provides a defensive physical barrier against pathogens in various plant species. Defense-induced lignification is a conserved basal defense mechanism in the plant immune response that is used as a biochemical marker of an activated immune response. The NAC-MYB-based gene regulatory network (NAC-MYB-GRN) regulates lignin biosynthesis (Liu et al., 2018; Ohtani and Demura, 2019).

Although many recent studies have shed light on various aspects of the growth–defense tradeoff, much remains to be learned about how TFs help coordinate plant growth with the appropriate responses to dynamic environmental conditions. In the current study, knocking down *ZmGT-3b* (encoding a GT-1 subfamily member) in young maize seedlings led to retarded growth, enhanced resistance to *F. graminearum* infection, and enhanced drought tolerance. ZmGT-3b positively regulates the expression of genes associated with photosynthesis (especially the critical seedling growth and light response regulator *ZmHY5*) and negatively regulates genes involved in plant defense responses.

This report shows that both photosynthesis- and defense-related gene expression is simultaneously regulated by the GT TF ZmGT-3b. This TF calibrates the plant growth–defense balance to coordinate metabolism during growth-to-defense transitions by optimizing the temporal and spatial expression of photosynthesis- and defense-related genes. ZmGT-3b might serve as a molecular hub connecting developmental/environmental signaling and secondary metabolite biosynthesis by repressing or activating specific pathways.

## RESULTS

### The Trihelix TF gene *ZmGT-3b* Responds Rapidly to *Fusarium graminearum* Infection in Maize Seedlings

We previously identified the quantitative trait locus (QTL) *qRfg1* on chromosome 10, which explained 36.6% of the total variation in maize resistance to *F. graminearum*-induced stalk rot (Yang et al., 2010). We then inoculated two maize near isogenic lines (NILs) carrying either the resistant *qRfg1* allele (R-NIL) or the susceptible *qRfg1* allele (S-NIL) with *F. graminearum* to evaluate the role of *qRfg1* in resistance to this pathogen (Ye et al., 2013). During the investigation of maize stalk rot disease resistance mechanism with the NILs, we found that the trihelix TF gene *ZmGT-3b* (*GRMZM2G325038*) was expressed at high levels in normal-especial R-NIL-seedling roots, but its expression was significantly decreased after inoculation (Supplemental Fig. S1A). Subsequently, we also detected a significant reduction in *ZmGT-3b* expression in response to *F. graminearum* inoculation in maize seedlings with or without the QTL *qRfg2* on chromosome 1 (Ye et al., 2019). ZmGT-3b belongs to the GT-1 clade of the plant-specific trihelix TF family. Based on transcriptome data, *ZmGT-3b* is primarily expressed in a few tissues, such as the primary root, ear primordium (2–8 µm), and embryo at 20 d after pollination (DAP) as well as the presheath and other tissues (Supplemental Fig. S1, B-C).

Consistently, the promoter region of *ZmGT-3b* (1500bp, 5’-upstream sequence of the starting codon ATG) contains 19 light-responsive *cis*-elements (LREs) and 21 defense or stress response-related *cis*-elements, including one W-box/Box-w1 (fungal elicitor responsive element or pathogen-inducible *cis*-element), two TC-rich repeats (*cis*-acting element involved in defense and stress responses), two MBSs (MYB binding site involved in drought inducibility), two ABREs (*cis*-acting element involved in the ABA response), eight TGACG-motifs (*cis*-acting regulatory element involved in the MeJA response), etc (Supplemental Table. S1). These elements/motifs likely allow the induction of *ZmGT-3b* in response to various development- and biotic/abiotic stress-related signals.

### *ZmGT-3b* Expression is Light-responsive, and ZmGT-3b Localizes to the Nucleus

We grew seedlings of maize inbred line B73-329 (wild type, used as the control [CK]) in the dark for 5 days and transferred them to light for 1 h and 2 h to analyze light-responsive *ZmGT-3b* expression. In agreement with the discovery of 19 LREs in the promoter of *ZmGT-3b*, *ZmGT-3b* expression was rapidly induced by light. This expression pattern is similar to the light-responsive expression of *ZmLHC117* and *ZmLHCII*, which encode components of the light harvesting complex (Fig. 1A). According to the B73 reference genome RefGen V4.32, *ZmGT-3b* contains three exons and encodes a predicted protein of 246 amino acids in length, containing the nuclear localization sequence (NLS) RKKLKRP at its N terminus. We obtained the full coding sequence of *ZmGT-3b* by RT-PCR and fused it to the N terminus of *GFP* under the control of the cauliflower mosaic virus (CAMV) 35S promoter. We transferred the resulting vector (*p35S*:*:ZmGT-3b-GFP*) into onion epidermal cells and *Nicotiana benthamiana* leaves by Agrobacterium (*Agrobacterium tumefasciens*)-mediated transient transformation and analyzed the subcellular localization of ZmGT-3b-GFP by confocal laser-scanning microscopy (using 488 nm excitation). ZmGT-3b-GFP localized specifically to the nuclei of epidermal cells (Fig. 1B).

**Figure 1.**
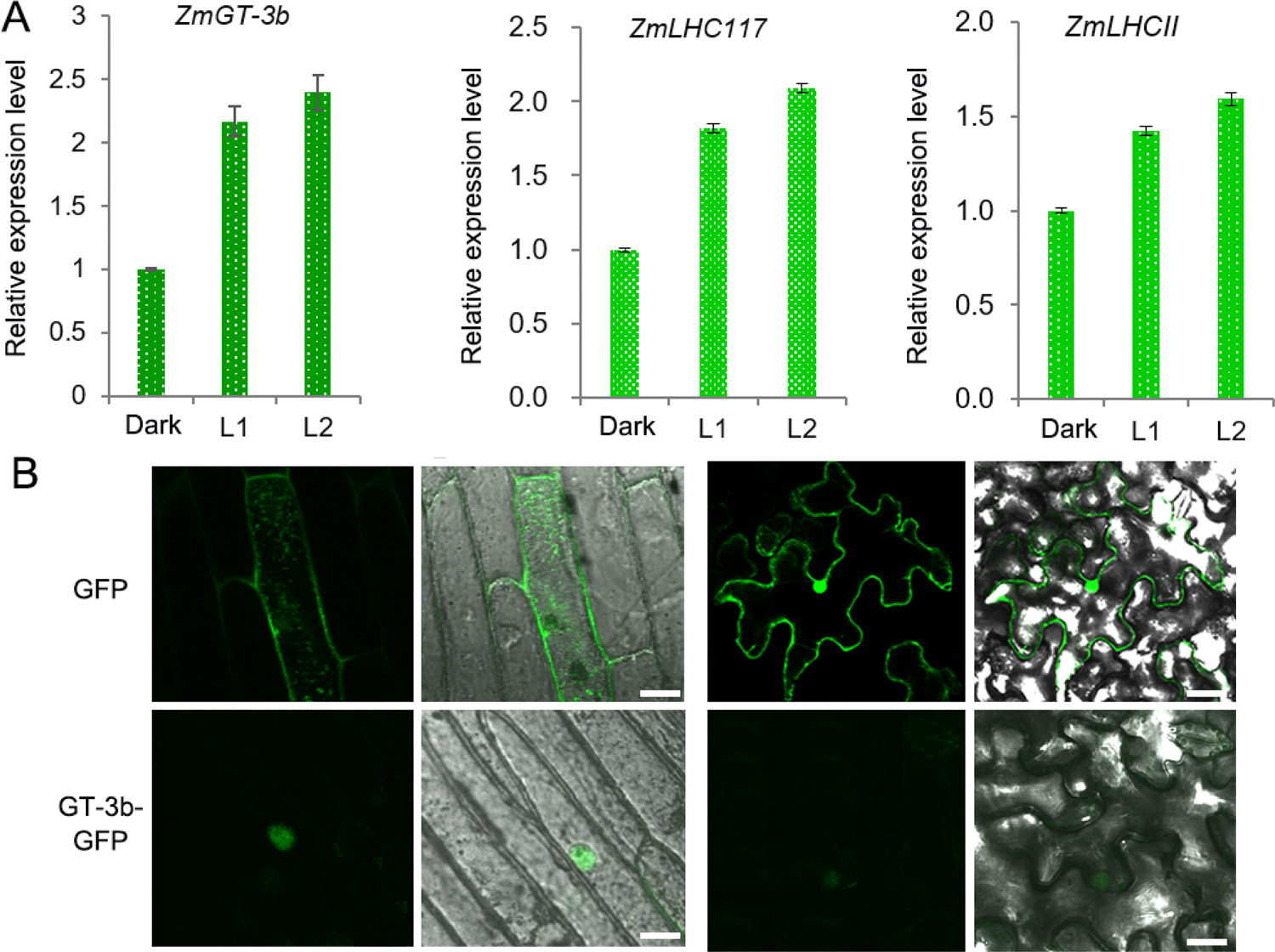
Light-responsive *ZmGT-3b* expression and subcellular localization of ZmGT-3b. A, The expression of *ZmGT-3b*, *ZmLHC117*, and *ZmLHCII* is induced by light. ‘Dark’ represents control seedlings (CK, maize inbred line B73-329, wild type) that were grown in the dark for 5 days; L1 and L2 represent CK seedlings transferred to the light for 1 h and 2 h after 5 days of growth in the dark, respectively. Error bars denote the mean ± SD of *n* = 3 replicates. B, ZmGT-3b-GFP localizes to the nucleus in onion epidermal cells and *Nicotiana benthamiana* leaves that were infiltrated with *Agrobacterium tumefaciens* carrying *p35S*: *ZmGT-3b-GFP* and observed 48 h after infiltration. GFP is the control transformed with empty vector containing the *p35S::GFP* expression cassette in the pCAMBIA1300 backbone. The images were taken under a confocal microscope. Bar =10 µm.

### Knockdown of *ZmGT-3b* Diminishes Seedling Growth by Reducing Photosynthetic Activity in Maize Seedling

We obtained mutant lines of *ZmGT-3b* by transforming a binary construct, which contained a cDNA fragment encoding the c-terminal 149aa of ZmGT-3b under the control of the maize *Ubiquitin* promoter, into maize receptor inbred line B73-329. Six transgenic events harboring the construct were obtained, self-crossed, and their homogenous seeds were harvested for functional analysis of *ZmGT-3b* in maize. Compared with CK seedlings, *ZmGT-3b* transcript levels in the primary roots of transgenic seedlings from four events (G3, G4, G6, and G7) were significantly reduced; the relative transcript level of G6 was only 0.11, and that of G7 was 0.375 compared with CK (Fig. 2A). We named the *ZmGT-3b* mutants with significantly reduced *ZmGT-3b* transcript levels *GT-KD*.

**Figure 2.**
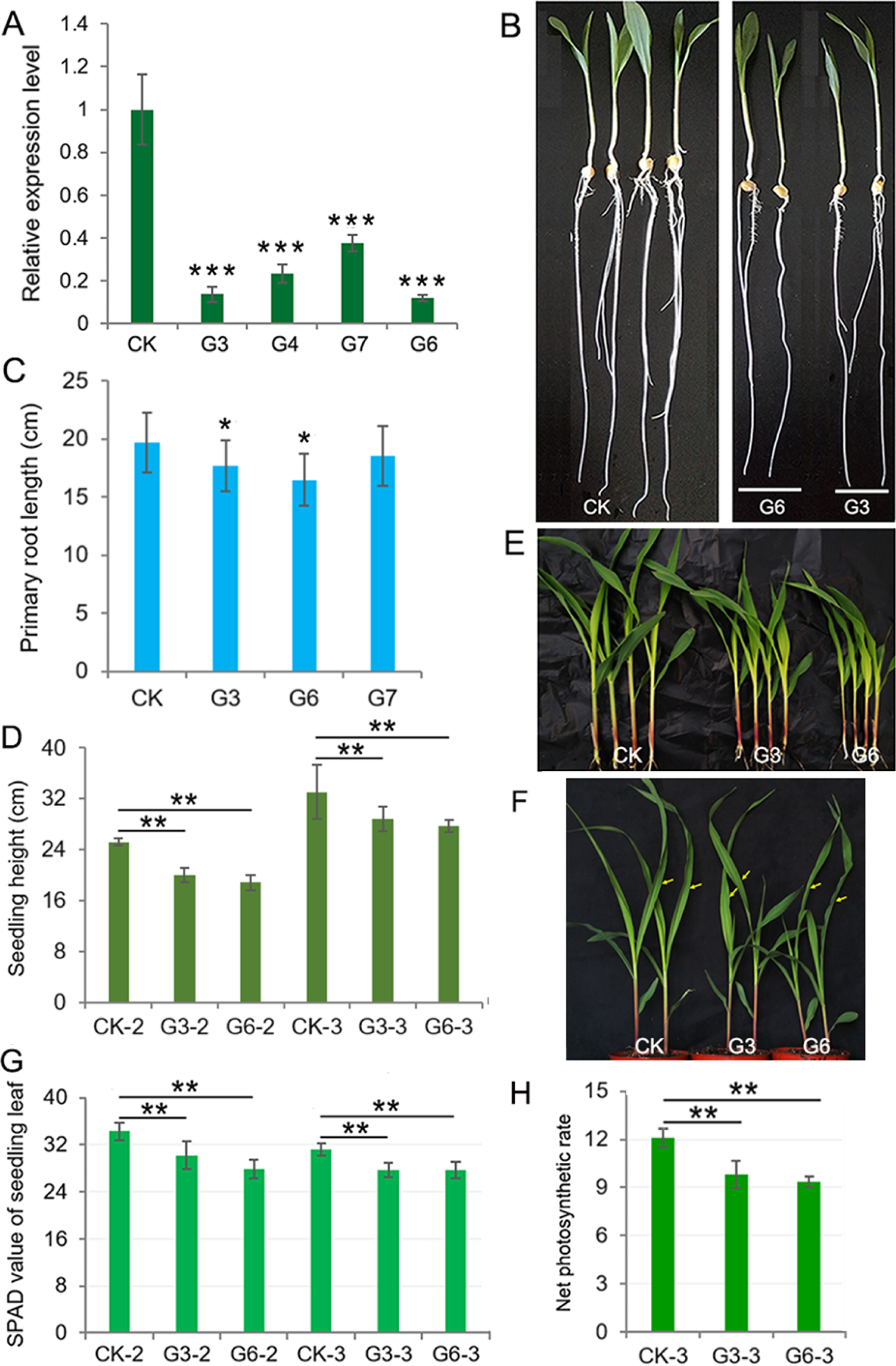
Significantly decreasing *ZmGT-3b* expression suppresses the growth of maize seedlings owing to reduced photosynthetic rates and chlorophyll contents. A, *ZmGT-3b* expression in the primary roots of transgenic maize seedlings is significantly reduced compared with CK (maize inbred line B73-329, wild type) at 7 days after germination (DAG). G3, G4, G6, and G7 represent seedlings from four transgenic events in which maize was transformed with the partial coding sequence of the C-terminal part of ZmGT-3b. Seedling phenotypes (B) and average primary root lengths (C) of 7-DAG maize seedlings. The average seedling height (D) and shoot phenotype (E for 12-DAG and F for 15-DAG) of soil-cultured maize seedlings grown in a greenhouse under natural light. Twelve-DAG seedlings with two leaves and a heart leaf are denoted as CK-2, G3-2, and G6-2, and 15-DAG seedlings with three leaves and a heart leaf are denoted as CK-3, G3-3, and G6-3 (used to measure net photosynthetic rate). The SPAD values (G) of the above seedlings were obtained from the central widest part of the newest expanded leaf, that is, the second leaf of 12-DAG seedlings or the third leaf of 15-DAG seedlings. (H) The net photosynthetic rate (μmol CO₂·m^−2^·s^−1^) of the central widest part of the third leaf of 15-DAG seedlings (indicated by an arrow in (F)) was measured from 09:00 to 12:00 under natural light in a greenhouse. Error bars denote the mean ± SD of three biological replicates. The asterisk indicates a statistically significant difference between CK and the *GT-KD* lines, as calculated by a paired Student’s t-test (* at *P* < 0.05, ** at *P* < 0.01, *** at *P* < 0.001).

The average primary root length of CK seedlings was ∼19.66 cm at 7 days after germination (7-DAG), the average primary root length of *GT-KD* seedlings was reduced by ∼10.25%, ∼16.17% and ∼5.72% for G3, G6 and G7, respectively (Fig. 2, B-C). The average seedling height was also significantly lower in G3 and G6 than in CK seedlings at both 12-DAG and 15-DAG when grown in a greenhouse under a 16h light/8h dark cycle, with reductions of ∼18.06% and ∼19.13% for G3 and G6 at 12-DAG, respectively (Fig. 2, D-E). When plants were grown in the field, plant height and 100-kernal-weight (HKW) of mature *GT-KD* plants did not significantly differ from those of CK plants grown in the same field (Supplemental Fig. S2). The retarded growth of *GT-KD* seedlings induced by the severe knockdown of *ZmGT-3b* suggests that *ZmGT-3b* is involved in positive regulation of maize seedlings root and shoot growth but has little effect on mature plant growth.

As trihelix TFs were initially found to participate in the light response, we examined the chlorophyll content and net photosynthetic rate (Pn) of *GT-KD* seedlings. The SPAD value is commonly used to estimate chlorophyll content. We measured the SPAD value at the center of the widest part of the newest expanded leaf of each seedling. The average SPAD value of 12-DAG CK seedlings was 34.3, while the average SPAD value of 12-DAG *GT-KD* seedlings decreased by ∼11.95% and ∼18.6% in G3 and G6, respectively, compared with CK. In 15-DAG seedlings, the average SPAD value decreased by ∼10.9% and ∼11.2% in G3 and G6, respectively, compared with CK. We also measured Pn from 09:00 to 12:00 in the center of the widest part of the newest expanded leaf of each 15-DAG seedling. The Pn value ranged from 11.3 to 12.7 μmol CO₂·m^−2^·s^−1^ in CK and from 8.9 to 11.2 μmol CO₂·m^−2^·s^−1^ in *GT-KD* seedlings. The average Pn decreased by ∼18.9% and ∼22.8% in G3 and G6, respectively compared with CK seedlings (Fig. 2, F-H). These results indicate that ZmGT-3b is involved in positive regulation of young seedling root and shoot growth by regulating chlorophyll biosynthesis and photosynthetic activity.

### *ZmGT-3b* Plays a Negative Role in Defense Responses and Drought Tolerance in Maize Seedlings

Both plant growth and immune responses consume large amounts of energy. The deployment of defense mechanisms is crucial for plant survival, thus the allocation of energy to defense activation generally comes at the expense of plant growth due to limited resources (Huot et al., 2014). Consistently, *ZmGT-3b* expression was significantly decreased in young CK seedlings after *F. graminearum* inoculation (Fig. 3A). *ZmGT-3b* knockdown led to retarded growth in young seedlings (Fig. 2, A–E), but improved resistance to *F. graminearum* infection. Both the shoot and root growth phenotypes of *GT-KD* seedlings were much better than CK seedlings following inoculation, and the disease severity index (DSI) of inoculated G3 and G6 seedlings was also markedly lower than that of inoculated CK seedlings (Fig. 3, B-C). However, following field inoculation of mature maize plants, the DSI of *GT-KD* plants was similar to (or higher than) that of CK plants (Fig. 3, D-E). Taken together, the finding that *ZmGT-3b* is only highly expressed in a few young tissues (Supplemental Fig. S1B-C), and that young *GT-KD* seedlings show retarded growth and improved disease resistance, suggests that *ZmGT-3b* is a positive regulator of growth and a negative regulator of disease resistance in maize seedlings.

**Figure 3.**
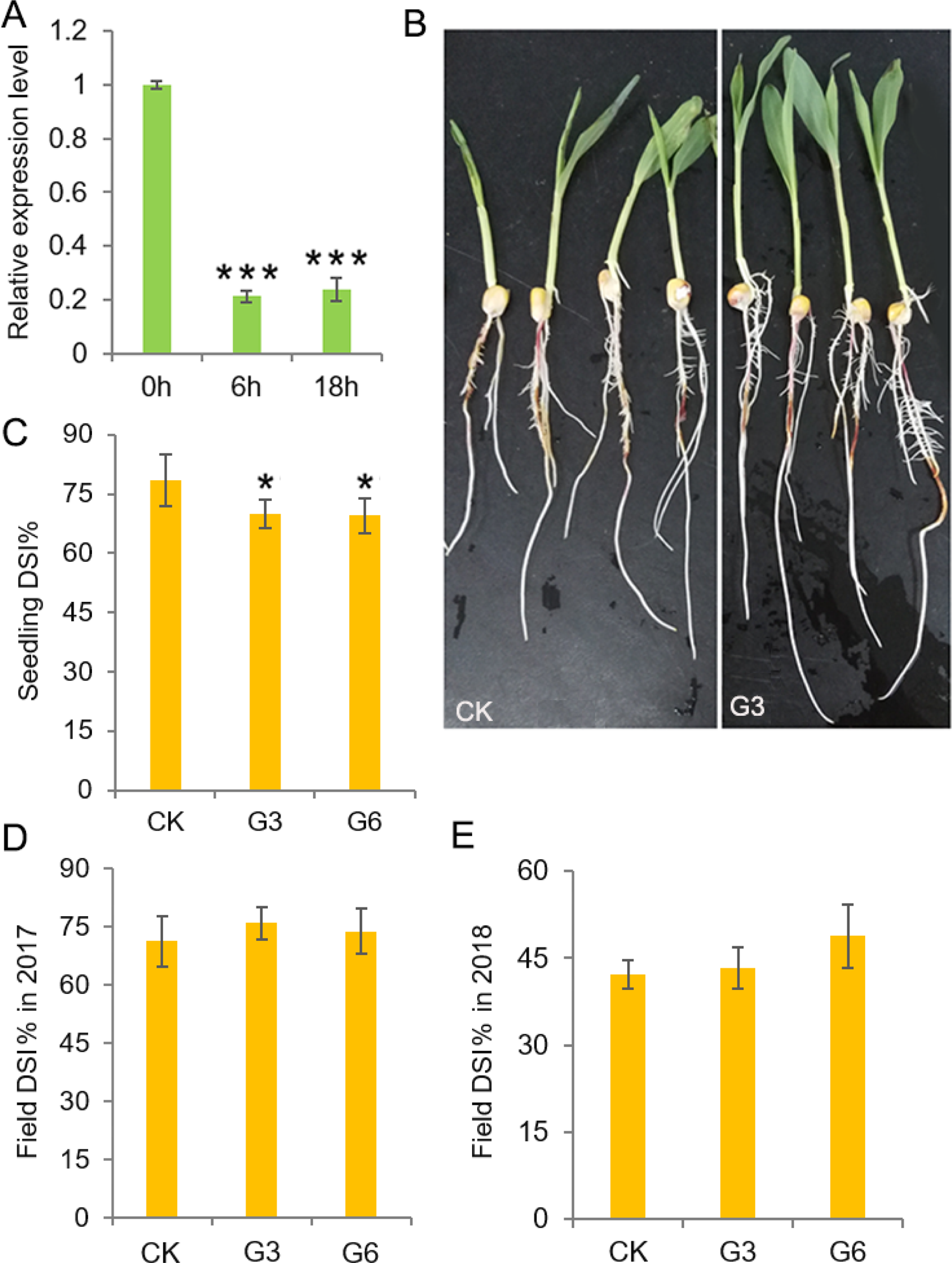
Analysis of the disease resistance of transgenic maize with significantly reduced *ZmGT-3b* expression. A, *ZmGT-3b* expression in young CK (maize inbred line B73-329, wild type) seedlings significantly decreased after *Fusarium graminearum* inoculation. ‘0h’, ‘6h’, and ‘18h’ indicate hours after inoculation of seedlings at 5 days after germination (DAG). Both shoot and root growth (B) were much better in inoculated *GT-KD* seedlings compared with inoculated CK seedlings with similar levels of disease severity, and the disease severity index (DSI) values of G3 and G6 seedlings were also significantly lower than that of CK seedlings (C). (D, E) The DSI of mature transgenic maize (*Zea mays*) plants was similar to (or higher than) that of CK plants in field inoculation experiments. Data of DSI are mean ± SD (*n* > 25 individual plants). The asterisk indicates a statistically significant difference between CK and the *GT-KD* lines, as calculated by a paired Student’s t test (* at *P* < 0.05, *** at *P* < 0.001).

Unexpectedly, when we stopped watering the growing seedlings after two-leaf stage, all CK seedlings severely wilted at 25-DAG, while *GT-KD* seedlings wilted less (see G7 seedlings in Fig. 4B), indicating that *GT-KD* seedlings were drought tolerant. Transgenic G3 and G6 seedlings were significantly shorter, but G7 seedlings were not significantly shorter, compared with CK (Fig. 4A). We then tested the leaf water loss rate (WLR) and survival rate (SR) of G3 and G7 seedlings (the growth of G7 seedlings was similar to CK under normal watering conditions). The leaf WLR and SR of both lines were significantly lower than those of CK (Fig. 4, C-D). We estimated the transpiration rate (TR) at the center of the widest part of the newest expanded leaf of each 15-DAG seedling (with three leaves and a heart leaf) (Fig. 4F) from 09:00 to 12:00. The estimated TR range of CK seedlings was 0.884-0.96 µmol H_2_O m^−2^ s^−1^, while that of *GT-KD* seedlings was 0.664-0.818 µmol H_2_O m^−2^ s^−1^. The average TR of G7 and G3 seedlings was ∼22.12% and ∼25.73% less than that of CK seedlings, respectively (Fig. 4E). These results indicate that *GT-KD* seedlings with knocked down *ZmGT-3b* expression were more drought tolerant than CK seedlings.

**Figure 4.**
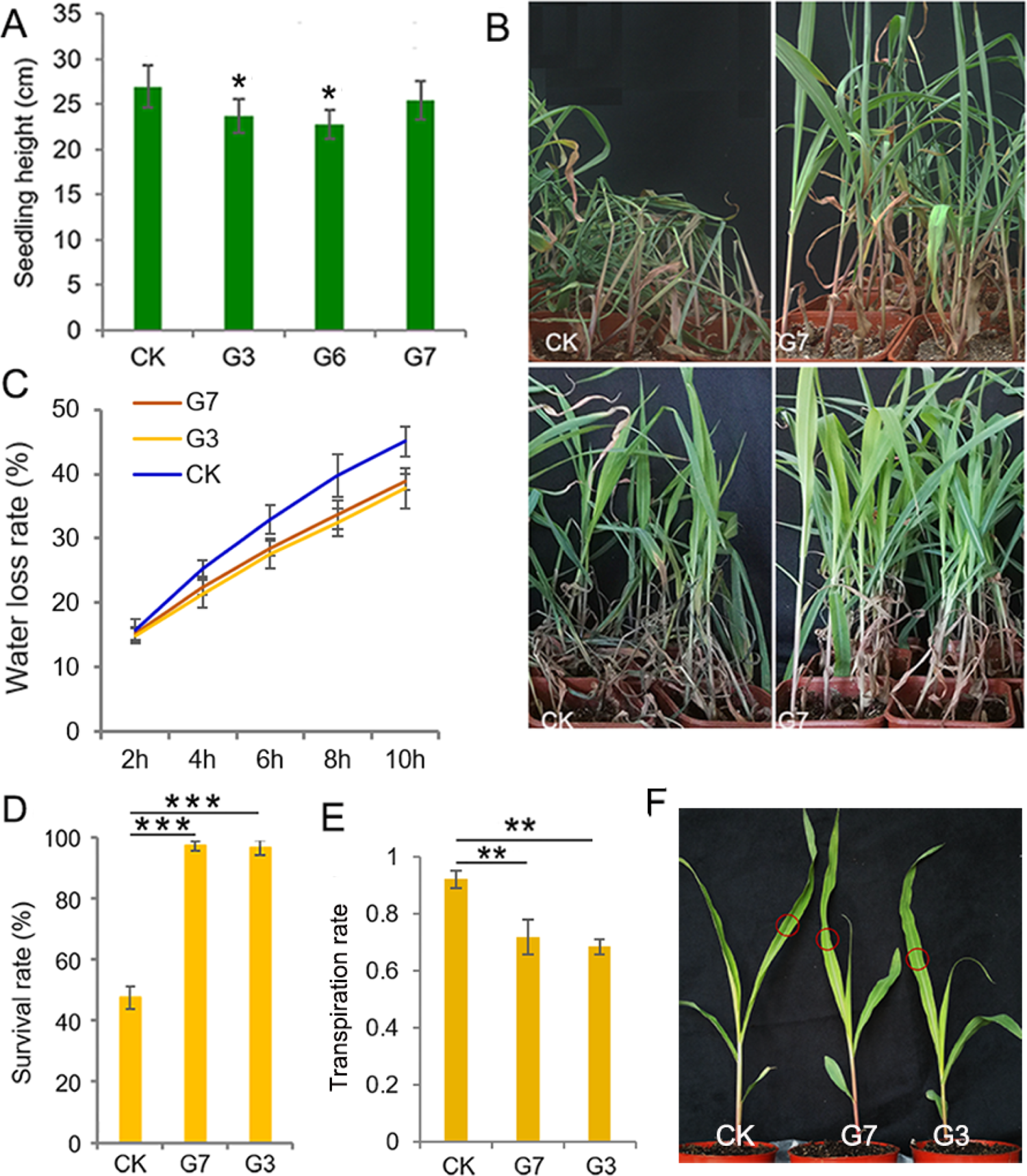
*ZmGT-3b* knockdown increases drought tolerance in *GT-KD* seedlings. A, Average height of 12-days after germination (DAG) seedlings grown in a growth chamber. Data are mean ± SD (*n* > 15 individual plants). B, Wilting (upper) and survival (bottom) of CK (maize inbred line B73-329, wild type) and G7 seedlings. Water was withheld from growing seedlings at the two-leaf stage, and the plants were re-watered when all CK seedlings were severely wilted (at 25-DAG). C, Leaf water loss rates of young maize (*Zea mays*) seedlings. The leaf water loss rates are shown as the means of the percentage of leaf water loss ± SD (*n* = 5). Three independent experiments were performed. (D–F) The survival rate (D), transpiration rate (TR, E), and seedling leaves (F) used for TR measurements. The estimated TR was obtained from the central widest part of the third leaf, which is the newest expanded leaf of 15-DAG CK or *GT-KD* seedlings (with three leaves and a heart leaf). The TR value was measured from 09:00 to 12:00 under natural light in a greenhouse, TR values are the mean ± SD (*n*=6). The asterisks indicate a statistically significant difference between CK and the *GT-KD* lines, as calculated by a paired Student’s t test (* at *p* < 0.05, ** at *P* < 0.01 and *** at *P* < 0.001).

### Photosynthesis-related Genes are Downregulated in the *ZmGT-3b* Knockdown Lines

To investigate the biological processes and genes regulated by ZmGT-3b, RNA-seq was used to compare the transcriptomes of 7-DAG seedlings of *GT-KD* (referred to as GT) with CK. The correlation coefficient (R) for the expression profiles of all transcripts between GT and CK was 0.87, suggesting that knockdown of *ZmGT-3b* affects overall gene transcription. We compared the transcriptional responses of *GT-KD* with CK seedlings and identified 950 differentially expressed genes (DEGs; upregulated ≥ 2-fold or downregulated ≤ 2-fold; *P*< 0.05), including 787 (83.7%) upregulated and 163 downregulated genes in *GT-KD* seedlings (Fig. 5A). The downregulated genes showed log2 fold change values of −6.69 to −1.01 and the upregulated genes showed log2 fold change values of 1.02 to 4.66. Of the downregulated DEG encoding proteins, 36 encode cellular components located in photosystems, 20 in plastoglobules, 41 in photosynthetic membranes, and 37 in the plastid thylakoid. Gene Ontology (GO) and Kyoto Encyclopedia of Genes and Genomes (KEGG) analysis revealed that the downregulated DEGs were significantly enriched in photosynthesis-related functional categories, such as photosynthesis (44 DEGs), photosynthesis light reaction (30 DEGs), photosynthesis light harvesting (17 DEGs), and photosynthesis antenna proteins (16 DEGs). This suggests that ZmGT-3b is a positive regulator of photosynthesis-related processes (Fig. 5, C-D; Supplemental Fig. S3). Therefore, the retarded growth of *GT-KD* seedlings might be caused by the transcriptional repression of growth-promoting (photosynthesis-related) genes caused by *ZmGT-3b* knockdown.

**Figure 5.**
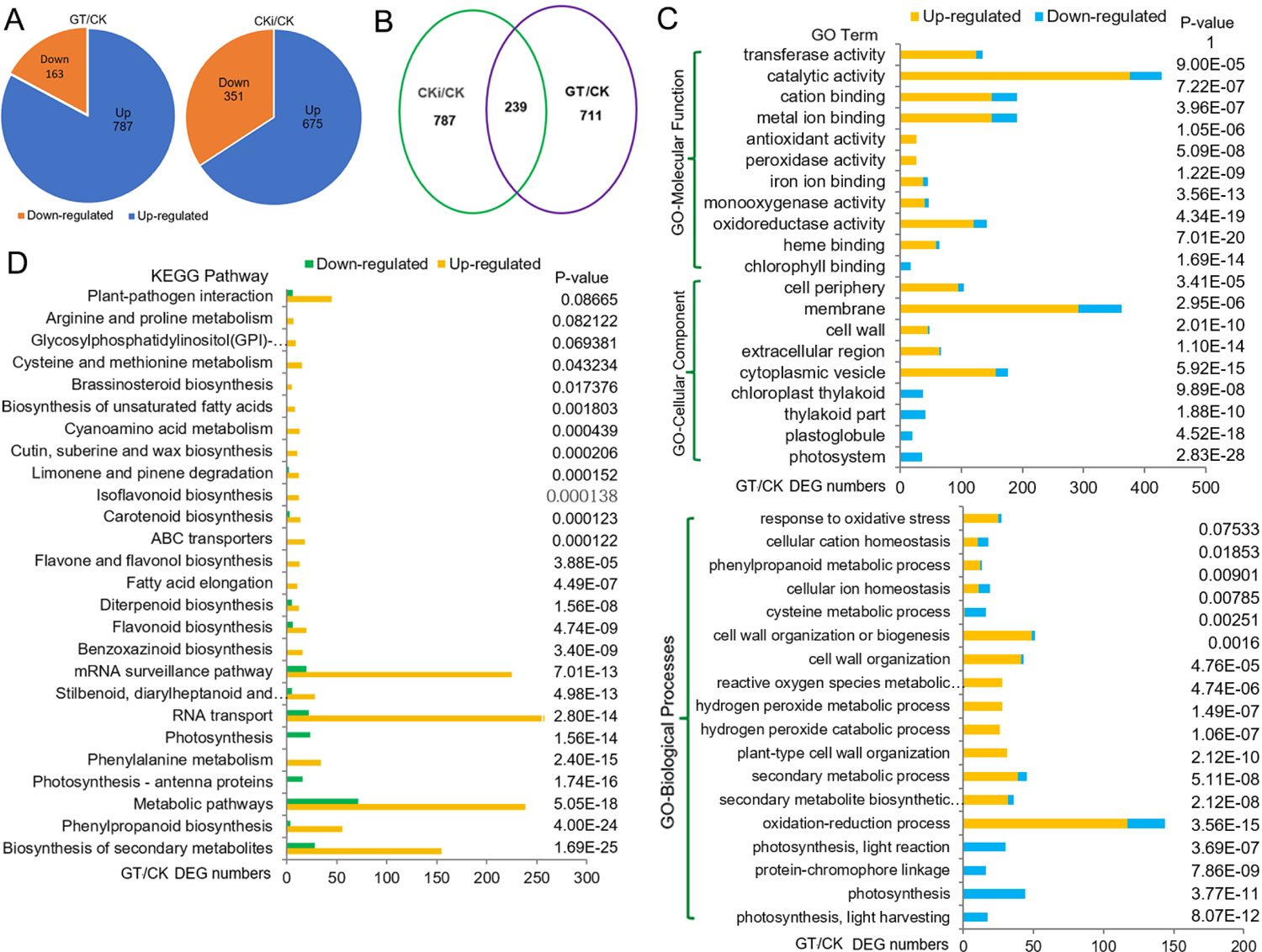
Transcriptome reprogramming induced by the knockdown of *ZmGT-3b*. A, Compared with CK (maize inbred line B73-329, wild type), *ZmGT-3b* knockdown induced the differential expression of 950 genes (GT/CK), while inoculation with *Fusarium graminearum* induced the differential expression of 1,026 genes (CKi/CK). GT represents seedlings from G3 and G6 transgenic plants grown under normal conditions without inoculation, CK and CKi represent control seedlings without or with inoculation, respectively. B, Two-hundred and thirty-nine differentially expressed genes (DEGs) were overlapping between CKi/CK and GT/CK. (C, D) Gene Ontology (GO) and Kyoto Encyclopedia of Genes and Genomes (KEGG) pathway functional analysis of the DEGs from GT/CK. Photosynthesis-related processes were reduced and defense response-related biological processes were enhanced by *ZmGT-3b* knockdown.

Among the significantly downregulated genes induced by *ZmGT-3b* knockdown, the bZIP TF gene *ZmHY5* (*ELONGATED HYPOCOTYL5*) showed an identical expression profile to *ZmGT-3b*; this gene was significantly downregulated in non-inoculated *GT-KD* seedlings and upregulated in inoculated *GT-KD* seedlings (Fig. 6A). Consistently, seven GT1 CONSENSUS (S000198) sites are discovered within 1500bp upstream of the start codon ATG of *ZmHY5*, which is the conserved DNA binding site of GT factors (Supplemental Table S1). HY5 is a central regulator of seedling development and light responses that promotes photomorphogenesis and mediates the positive effect of shoot illumination on root growth (Chen et al, 2016; Gangappa and Botto, 2016). This TF modulates photosynthetic capacity by controlling the expression of chlorophyll biosynthesis- and photosynthesis-related genes (Kobayashi et al., 2012; Toledo-Ortiz et al., 2014).

**Figure 6.**
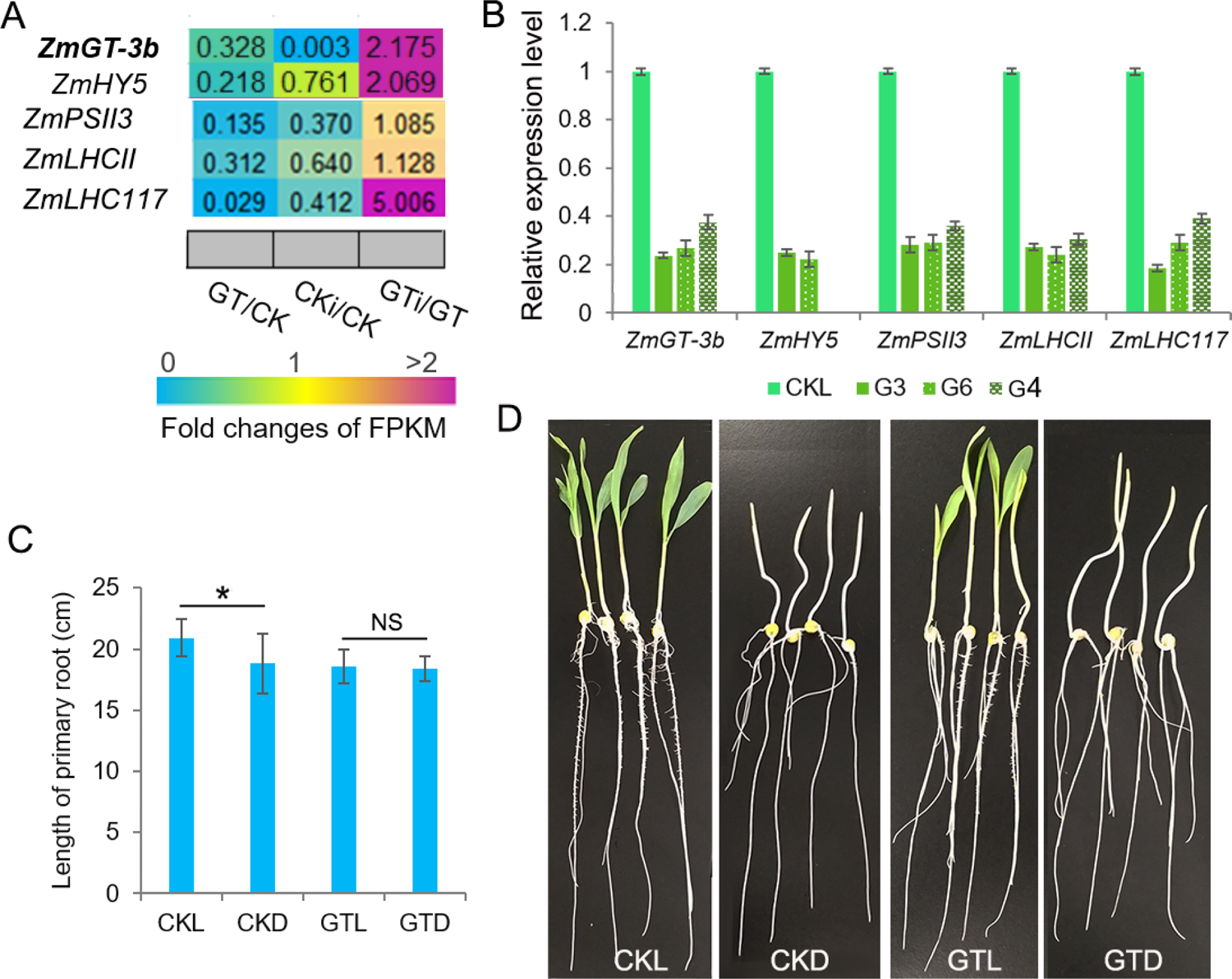
*ZmGT-3b* knockdown dramatically downregulates genes associated with photosynthesis-related functional categories. A, Relative expression (fold) of photosynthesis-related genes from *ZmGT-3b* knockdown (GT) and CK (B73-329) maize seedlings at 7 days after germination (DAG) with (CKi, GTi) or without inoculation (CK, GT) based on transcriptome sequencing. B, The expression of light-responsive genes, such as *ZmGT-3b, ZmHY5, ZmPSII3, ZmLHCII* and *ZmLHC117*, decreased in seedlings with knocked down *ZmGT-3b* expression, as confirmed by qRT-PCR. CK represents control seedlings (B73-329), G3, G6 and G4 represent seedlings from different transgenic events with knocked down *ZmGT-3b* expression. All seedlings were grown under normal conditions and sampled at 7-DAG. (C, D) *GT-KD* seedlings with significantly reduced *ZmHY5* expression caused by *ZmGT-3b* knockdown showed disrupted shoot illumination-promoted root growth. The average primary root length of CK seedlings grown in the light was significantly higher than that of plants grown in the dark (C), while the average primary root lengths of *GT-KD* seedlings were similar in both the light and dark (D). CKL/CKD represent CK seedlings grown in the light/dark; GTL/GTD represent *GT-KD* seedlings grown in the light/dark. Values are the mean ± SD (*n* > 15). The asterisk * represents a significant difference at *P* < 0.05 (according to a paired Student’s t-test); NS, not significant.

To verify the RNA sequencing data, we performed qRT-PCR to compare gene expression levels between *GT-KD* and CK seedlings. *ZmGT-3b*, *ZmHY5*, and randomly selected photosynthesis-related genes such as *ZmPSII3, ZmLHCII*, and *ZmLHC117* were significantly downregulated in various *GT-KD* seedlings (Fig. 6B). Moreover, the primary roots of CK seedlings grown in the light were significantly longer than those grown in the dark, whereas the primary root lengths of *GT-KD* seedlings, which showed dramatically reduced *ZmGT-3b* and *ZmHY5* expression, were similar in plants grown in the light and in the dark (Fig. 6, C-D). These results indicate that the knockdown of *ZmGT-3b* led to reduced *ZmHY5* expression, which disrupted the promotion of root growth via shoot illumination in *GT-KD* seedlings. These findings are consistent with the reduced chlorophyll contents and photosynthetic rates of *GT-KD* seedlings (Fig. 2, D–H). These results suggest that ZmGT-3b regulates seedling growth by modulating chlorophyll biosynthesis and photosynthetic activity via the transcriptional regulation of photosynthesis-related genes.

### *ZmGT-3b* Knockdown Induces Defense-related Transcriptional Reprogramming without Immunity Activation

Most proteins encoded by the upregulated DEGs were associated with the membrane, cell periphery, or cytoplasmic vesicle, and with molecular functions including catalytic activity, oxidoreductase activity, transferase activity, metal ion binding, cation binding, and iron ion binding. The upregulated DEGs were enriched in oxidation–reduction processes, secondary metabolic processes, plant-type cell wall organization, and reactive oxygen species (ROS) metabolic processes. These upregulated DEGs significantly contribute to the biosynthesis of secondary metabolites, especially the biosynthesis of phenylpropanoid, stilbenoid, diarylheptanoid, gingerol, benzoxazinoid, flavonoid, diterpenoid, flavone, flavonol, and carotenoid (Fig. 5, C-D). Almost all of these functional categories support basal defense responses to various biotic/abiotic stresses, suggesting that ZmGT-3b acts as a negative regulator of plant defense response-related biological processes.

Dramatic transcriptional reprogramming occurs during the induction of plant immune responses, allowing the plant to prioritize defense-over growth-related cellular functions. The correlation coefficient (R) for the expression profiles of all transcripts between the inoculated CK (CKi) and CK (CKi/CK) was 0.86, i.e., close to the value of 0.87 between GT and CK (GT/CK). This indicates that inoculation and *ZmGT-3b* knockdown (GT/CK) have similar effects on general gene transcription. Compared with untreated CK seedlings, inoculation induced 1,026 DEGs, including 239 DEGs that were simultaneously induced by *ZmGT-3b* knockdown. This indicates that overlapping events or defense signaling pathways control gene expression in response to inoculation (CKi/CK) or *ZmGT-3b* knockdown (GT/CK) (Fig. 5B).

The upregulated DEGs induced by inoculation (CKi/CK) were also significantly enriched in GO terms associated with biosynthesis of secondary metabolites, especially biosynthesis of phenylpropanoid, stilbenoid, diarylheptanoid, and gingerol, benzoxazinoid, flavonoid, diterpenoid, flavone and flavonol, carotenoid, phenylalanine metabolism and plant–pathogen interaction. These results are similar to the transcriptome reprogramming induced by *ZmGT-3b* knockdown (GT/CK), with many more DEGs enriched in photosynthesis, RNA transport, and mRNA surveillance pathway (Supplemental Fig. S4). Therefore, considerable commonalities were detected in the transcriptional reprogramming induced by *ZmGT-3b* knockdown or inoculation. *ZmGT-3b* knockdown induced defense-related transcriptional reprogramming without immunity activation to upregulate or activate basal defense-related genes. Therefore, ZmGT-3b might act as a transcriptional repressor of genes involved in basal defense responses.

Furthermore, 1,049 genes were differentially expressed in inoculated *GT-KD* versus inoculated CK seedlings (GTi/CKi) (Supplemental Fig. S5A). Of these, 254 DEGs were shared between the inoculated transcriptome pair GTi/CKi and the non-inoculated transcriptome pair GT/CK (Supplemental Fig. S5B). The most highly enriched GO functional categories were similar, and most highly enriched KEGG pathways were shared between the two transcriptome pairs. The transcriptional differences between the non-inoculated GT/CK transcriptome emphasized in downregulated photosynthesis-related processes. Most of the shared functional categories are involved in basal defense responses to various biotic/abiotic stresses (Supplemental Fig. S5, C-D), pointing to considerable commonalities in their transcriptional responses.

However, the R for the expression profiles of all transcripts between CKi and CK was 0.86, and that between GTi and GT was 0.63, indicating that inoculation affected overall gene transcription to different degrees in the two genotypes. Compared with non-inoculated seedlings, inoculation induced the expression of 1,026 and 1,340 DEGs in CK and *GT-KD*, respectively, with 462 genes induced in both genotypes (Supplemental Fig. S5, C-D). This suggests that overlapping signaling pathways control gene expression in these two types of seedlings in response to inoculation. The transcriptional reprogramming was much stronger between inoculated and non-inoculated *GT-KD* seedlings (GTi/ GT) than between inoculated and non-inoculated CK seedlings (CKi/CK). Many more DEGs were enriched in almost all top defense-related GO and KEGG functional categories, except for biosynthesis of brassinosteroids and zeatin, which had more DEGs enriched in CKi/CK (Supplemental Fig. S5, E-F). These results are consistent with the notion that inoculated *GT-KD* have better disease resistance and lower DSI than CK seedlings.

### *ZmGT-3b* Knockdown Leads to the Upregulation of Defense-related Genes

*ZmGT-3b* knockdown dramatically upregulated genes that function in defense responses to various biotic/abiotic stress, including genes encoding six PR proteins, three bZIPs, 18 MYBs, six WRKYs, eight NACs, eight ethylene-response factors (ERFs, the largest group of the AP2/EREBP family), and three bHLHs, which were all significantly upregulated in *GT-KD* versus CK seedlings (Supplemental Fig. S6). Many members of these TF families reportedly function in plant responses to various biotic or abiotic stresses. Members of the MYB, ERF, bZIP, and NAC TF families are well-known transcriptional regulators of genes required for drought tolerance, and MYBs are important regulators of both secondary cell wall biosynthesis and abiotic stress tolerance (Mizoi et al., 2012; Baldoni et al., 2015). Among the TF genes upregulated by *ZmGT-3b* knockdown, some were significantly upregulated in *GT-KD* seedlings without inoculation and in CK seedlings with inoculation, contrasting to *ZmGT-3b*, including *ZmWRKY11*, *ZmWRKY69*, *ZmMYB36*, *ZmMYB93*, *ZmIBH1* (a basic helix-loop-helix, bHLH)*, ZmbHLH28*, *ZmNAC67, ZmMYB8*, *ZmbZIP53*, *ZmbZIP7*, and *ZmbZIP8* (Fig. 7A). The upregulated expression of these TF genes and a few well-known defense-related genes such as *ZmDRR206*, *ZmPR1*, and *ZmPR-STH21* was confirmed by qRT-PCR in different *GT-KD* seedlings with knocked down *ZmGT-3b* expression compared with CK seedlings (Fig. 7B). Therefore, various TF genes and defense-related genes were upregulated by *ZmGT-3b* knockdown.

**Figure 7.**
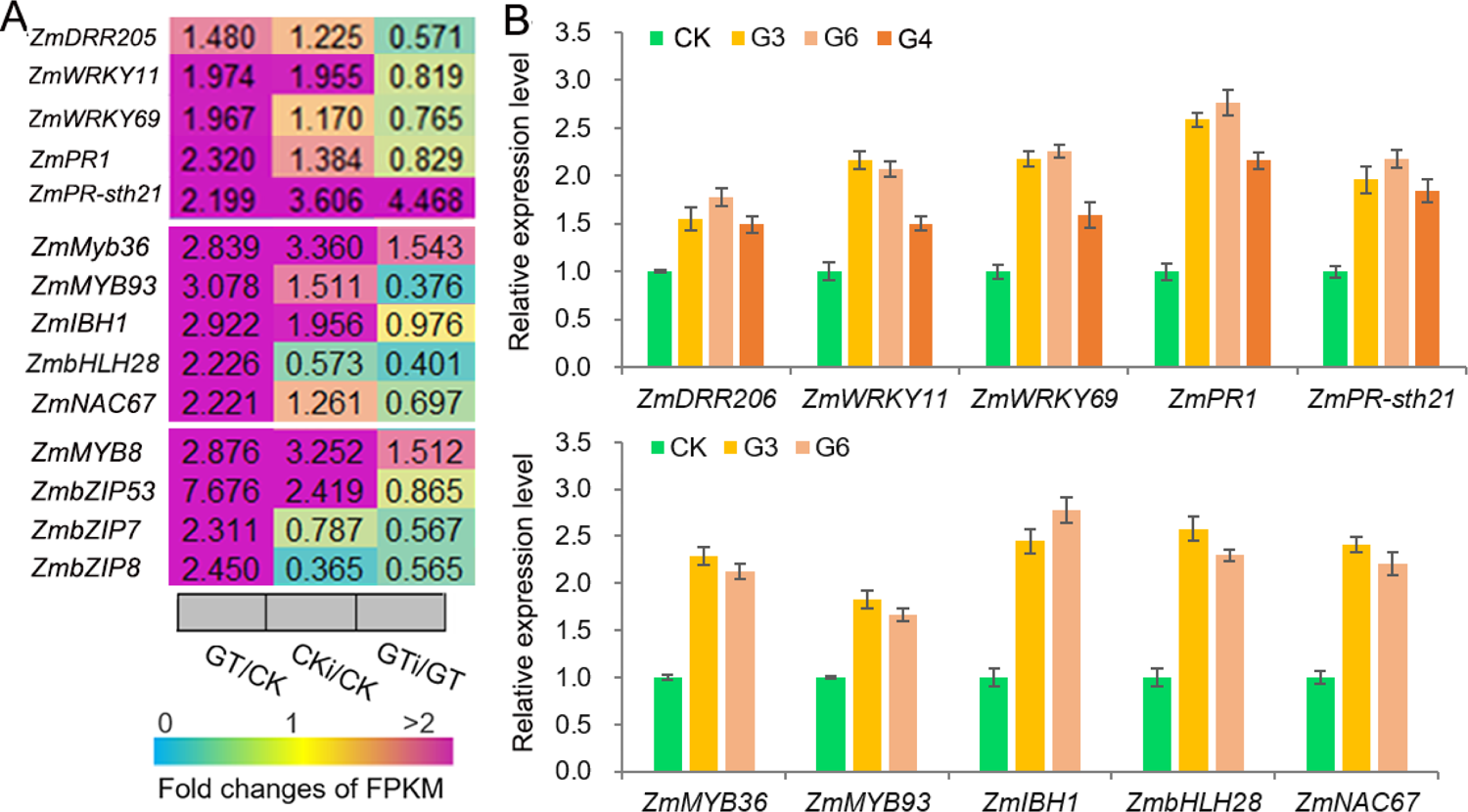
*ZmGT-3b* knockdown dramatically upregulates genes encoding multiple transcription factors (TFs) involved in plant defense response to various biotic/abiotic stresses. A, Relative expression levels (fold) of TF genes, including MYBs, bZIPs, NACs, bHLHs genes, as revealed by transcriptome sequencing of *ZmGT-3b* knockdown (GT) and CK (maize inbred line B73-329, wild type) maize (*Zea mays*) seedlings at 7 days after germination (DAG) with (CKi, GTi) or without inoculation (CK, GT). B, Defense-related genes such as *ZmDRR206*, *ZmPR1*, and *ZmPR-STH21* and multiple defense-related TF genes such as *ZmWRKY11*, *ZmWRKY69*, *ZmMYB36*, *ZmMYB93*, *ZmIBH1, ZmbHLH28* and *ZmNAC67* were upregulated in *GT-KD* seedlings, as confirmed by qRT-PCR. CK represents control seedlings (B73-329), and G3, G6, and G4 represent seedlings from different transgenic events with knocked down *ZmGT-3b* expression. All seedlings were grown under normal conditions and sampled at 7-DAG.

### *ZmGT-3b* Knockdown Increases the Biosynthesis of Cell Wall Components Independently of Immune Activation

Based on the analysis of transcriptome reprogramming induced by *ZmGT-3b* knockdown (GT/CK), many proteins encoded by the DEGs are involved in the biosynthesis of secondary metabolites, especially phenylpropanoid (Fig. 5, C-D), which is the first critical step for lignin biosynthesis. Lignin is one of the most important secondary metabolites and defense-induced lignin biosynthesis plays a major role in basal immunity. We therefore measured the contents of the major components of the plant cell wall, such as cellulose, semi-cellulose, and lignin in seedlings.

Compared with CK seedlings, the contents of cellulose (∼5.4% increase), semi-cellulose (∼7.3% increase), acid soluble lignin (ASL, ∼22.7% increase), and lignin (∼4.64% increase) were markedly higher in 12-DAG *GT-KD* maize seedlings, whereas the content of acid-insoluble lignin was not (AIL) (Fig. 8A). Arabinose levels are positively associated with lignin levels, and high concentrations of xylose are important in defense responses (Li et al., 2015). The levels of both arabinose (∼6.31% increase) and xylose (∼6.7% increase) were significantly higher in 12-DAG *GT-KD* than CK seedlings (Fig. 8B). Consistently, almost all genes encoding the critical enzymes in the lignin biosynthesis pathway were strongly upregulated in *ZmGT-3b* knockdown seedlings, including genes encoding three phenylalanine ammonia-lyases (PALs), two 4-coumarate CoA ligases (4CLs), six hydroxycinnamoyl-CoA shikimate/Quinate hydroxycinnamoyl transferases (HCTs), one caffeoyl-CoA *O*-methyltransferase (CCoAOMT), one cinnamoyl-CoA reductase (CCR), six caffeic acid *O*-methyltransferases (COMTs), three laccases (LACs), four dirigent proteins (DPs), and 23 peroxidases (PODs); genes encoding cinnamate 4-monooxygenase (C4M) and cinnamyl alcohol dehydrogenase (CAD) were not upregulated in these seedlings. Seven genes encoding Casparian strip membrane proteins (CASPs) were also significantly upregulated in 12-DAG *GT-KD* vs. CK seedlings (Fig. 8, C-D, Supplemental Fig. S6). Furthermore, all 22 cellulose-synthase (Cesa) genes with highly abundant transcripts (with FPKM>20) showed elevated expression levels in *GT-KD* seedlings, and four of these genes (with FPKM>50) had FC>1.5 in GT-KD versus CK seedlings (Fig. 8E). These data suggest that *ZmGT-3b* knockdown promoted secondary metabolite biosynthesis, especially lignin and cellulose biosynthesis, which occurred independently of immune activation in maize seedlings.

**Figure 8.**
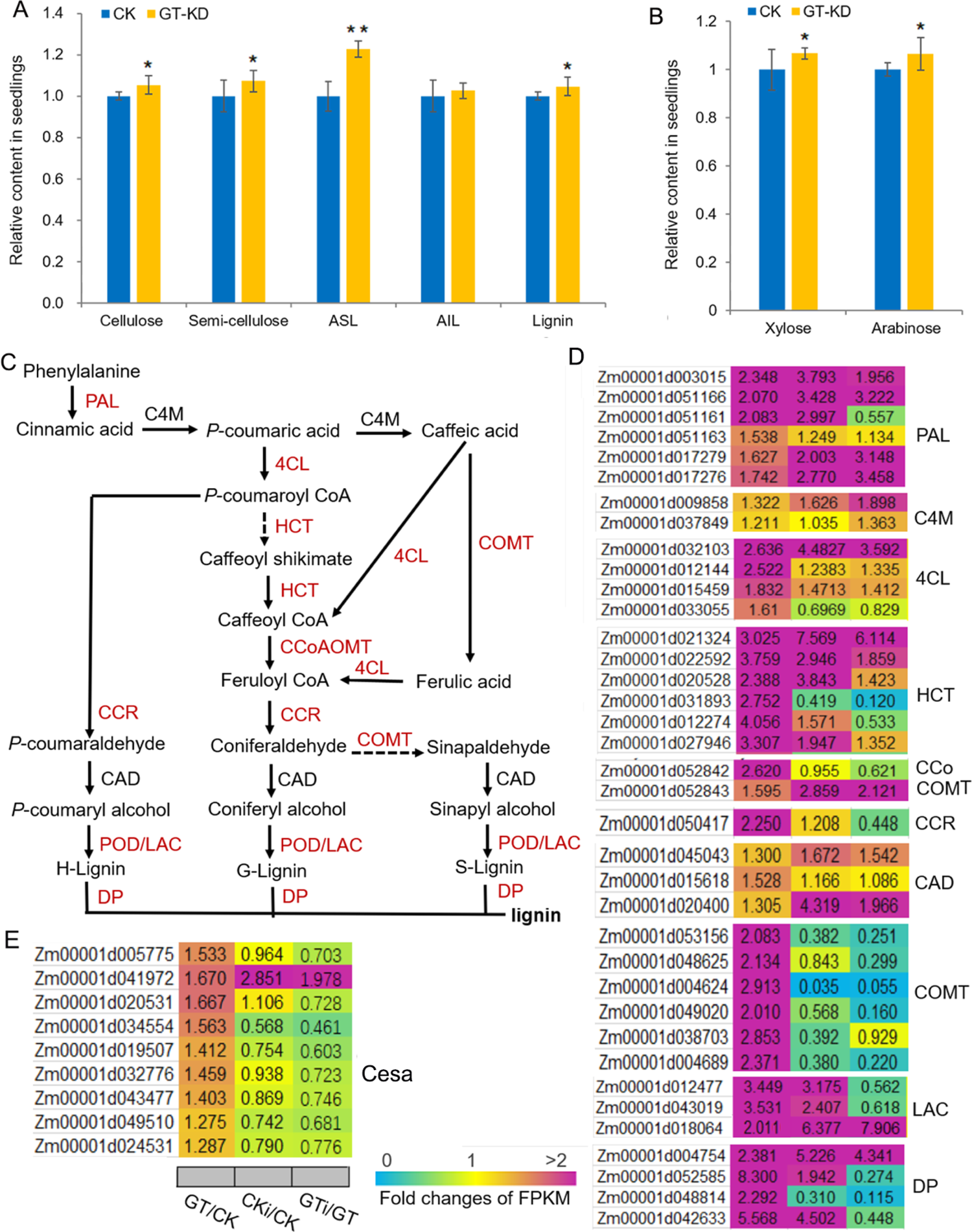
The contents of the major plant cell wall components and critical genes in the lignin biosynthesis pathway are upregulated in *ZmGT-3b* knockdown maize seedlings. (A) The cellulose, semi-cellulose, acid soluble lignin (ASL), and lignin contents were significantly higher in 12-days after germination (DAG) *GT-KD* versus CK (maize inbred line B73-329, wild type) seedlings, while the acid-insoluble lignin (AIL) contents were similar. (B) Both arabinose and xylose levels were significantly higher in *GT-KD* seedlings than CK seedlings. Values are the mean ± SD (*n* = 3). The asterisks represent a significant difference at * *P* < 0.05, ** *P* < 0.01 (according to a paired Student’s t-test). (C) The general lignin biosynthesis pathway, including the related enzymes. Red indicates significantly upregulated, and black indicates not significantly upregulated, in *GT-KD* seedlings. (D) Relative expression levels (fold) of genes involved in lignin biosynthesis (D) and cellulose synthase genes (E) in *GT-KD* seedlings. The data were obtained by transcriptome sequencing of *ZmGT-3b* knockdown (GT) and CK seedlings with (CKi, GTi) or without inoculation (CK, GT). PAL, phenylalanine ammonia-lyase; C4M, cinnamate 4-monooxygenase; 4CL, 4-coumarate CoA ligase; HCT, hydroxycinnamoyl-CoA shikimate/quinate hydroxycinnamoyl transferase; CCoAOMT, caffeoyl-CoA *O*-methyltransferase; CCR, cinnamoyl-CoA reductase; CAD, cinnamyl alcohol dehydrogenase; COMT, caffeic acid *O*-methyltransferase; LAC, laccase; POD, peroxidase; and DP, dirigent protein (DP).

Many genes upregulated in *ZmGT-3b* knockdown plants encode proteins involved in metal ion binding, cation binding, or iron ion binding (Fig. 5, C-D). Thus, we measured the contents of mineral elements in CK and *GT-KD* seedlings at 12-DAG. Compared with CK seedlings, potassium (K^+^), phosphorus (P), and copper (Cu) levels were significantly higher, while aluminum (Al) and iron (Fe) levels were significantly lower, in *GT-KD* seedlings.

However, Zn, Mg, and Na levels did not significantly differ in GT-KD vesus CK seedlings (Supplemental Fig. S7A). Consistent with the different contents of various mineral elements, genes encoding transporters of these elements showed considerably different transcript levels in GT-KD versus CK seedlings, including genes encoding phosphate, potassium, copper, vacuolar iron, and zinc transporters. Finally, two sulfate transporter genes were significantly upregulated in *GT-KD* versus CK seedlings (Supplemental Fig. S7B). These results suggest that *ZmGT-3b* knockdown altered cellular osmotic conditions by increasing/decreasing the contents of various mineral elements by affecting the take-up of inorganic ions from the environment using the corresponding transporters.

## DISCUSSION

The growth–defense tradeoff in plants is associated with the limited availability of resources, which requires the plant to prioritize growth or defense, depending on dynamic external and internal factors. This balance is important for agriculture and natural ecosystems due to the vital importance of these processes for plant survival, reproduction, and plant fitness (Huot et al., 2014). Numerous studies have revealed the crucial roles of TFs in regulating diverse plant growth, development, and biotic or abiotic stress responses, including the trihelix TFs (Kaplan-Levy et al., 2012). Here, we showed that *ZmGT-3b* transcript levels were significantly decreased during the interaction of maize seedlings with *F. graminearum*. In addition, *ZmGT-3b* knockdown in maize seedlings led to retarded growth, improved resistance to *F. graminearum*, and enhanced drought tolerance (Fig. 2–4). Moreover, *ZmGT-3b* is associated with the positive regulation of photosynthesis-related genes and the negative regulation of defense-related genes, as *ZmGT-3b* knockdown severely downregulated the expression of photosynthesis-related genes and significantly upregulated the expression of defense-related genes at the same time (Fig. 5-7, Supplemental Fig. S, 5-6). These findings suggest that ZmGT-3b involves in pathogen attack induced suppression of photosynthesis activity, and coordinates metabolism during growth-to-defense transitions by optimizing the temporal and spatial expression of photosynthesis- and defense-related genes, uncovering a molecular mechanism underlying the growth– defense tradeoff.

### ZmGT-3b Modulates Seedling Growth via the Transcriptional Regulation of Photosynthesis-related Genes

The conserved DNA-binding domains of plant-specific trihelix TFs are often found at the N terminus. By contrast, the C terminus, which harbors a hydrophilic region, is less conserved and probably acts as the activation domain (Qin et al., 2014; Kaplan-Levy et al., 2014). Trihelix TFs control the transcription of light-regulated genes as well as genes involved in plant development and stress responses (Kaplan-Levy et al., 2012). During these transcriptional regulatory processes, trihelix TFs form homodimers or heterodimers by participating in interactions with other classes of TFs (Xie et al., 2009; Li et al., 2015). Consistent with the light-inducible expression profile of *ZmGT-3b*, we identified 19 light-responsive elements in the promoter of this gene (Fig. 1A, Supplemental Table. S1).

We obtained *GT-KD* mutants with severely reduced *ZmGT-3b* levels by transforming maize inbred line B73-329 with a cDNA fragment encoding the C-terminal 149 aa of ZmGT-3b under the control of the maize *Ubiquitin* promoter. This mutant might produce protein without the conserved N-terminal DNA-binding domains, ultimately leading to *ZmGT-3b* knockdown. These plants exhibited diminished growth at the seedling stage, with shorter primary roots and reduced seedling height (Fig. 2, A–D). However, *ZmGT-3b* knockdown had no effect on the growth of mature maize plants in the field (Supplemental Fig. S2). The effect of *ZmGT-3b* knockdown on seedling growth might be associated with the expression profile of *ZmGT-3b* (primarily expressed in a few specific young tissues, such as primary roots and ear primordium) and the elements for meristem and root expression in its promoter (Supplemental Fig. S1, Table S1).

Light perception activates many TFs from various families, such as bZIP, bHLH, MYB, GATA, and GT1. These TFs bind to various LREs such as G, GT1, GATA, and MREs, leading to massive transcriptional reprogramming (Gangappa and Botto, 2014, 2016). Among these TFs, the role of HY5 as a master transcriptional regulator is conserved across plant species. HY5 mediates the light-responsive coupling of shoot growth and photosynthesis with root growth and nitrate uptake, and it functions as the center of a transcriptional network hub connecting different processes such as hormone, nutrient, abiotic stress (abscisic acid, salt, cold), and ROS signaling pathways (Chen et al., 2016; Gangappa and Botto 2016; Burman et al., 2018). HY5 activates its own expression and is a critical player in seedling development and responses to light by turning on or off many genes involved in fundamental developmental processes such as cell elongation, pigment accumulation (chlorophyll and anthocyanin), flowering, and root development (Kobayashi et al., 2012; Toledo-Ortiz et al., 2014; Chen et al. 2016). The activator or repressor activity of HY5 in transcription regulation during plant growth and light responses depends on its interacting partners, as HY5 does not have its own activation or repression domain. The primary function of HY5 in promoting transcription may depend on other, likely light-regulated, factors (Burko et al., 2020).

*ZmGT-3b* knockdown induced the downregulation of 163 DEGs in *GT-KD* seedlings. GO and KEGG analysis revealed that these downregulated DEGs were significantly enriched in the functional category photosystem, but not in carbon metabolism-related functional categories, including photosynthesis-light harvesting, -light reaction, protein-chromophore linkage (antenna proteins), and photosynthesis (Fig. 5, C-D, Supplemental Fig. S3). *ZmHY5* was significantly downregulated by *ZmGT-3b* knockdown, as confirmed by qRT-PCR in independent *GT-KD* lines (Fig. 6B). The significantly decreased /increased transcript levels of both *ZmHY5* and *ZmGT-3b* in *GT-KD* seedlings without/with inoculation (Fig. 6A), the strongly reduced chlorophyll content (SPAD value) and net photosynthetic rates of *GT-KD* seedlings (Fig. 2, G-H), and the disruption of the effect of shoot illumination on promoting root growth by *ZmGT-3b* knockdown-induced *ZmHY5* downregulation (as root lengths were similar in *GT-KD* seedlings grown in both the light and dark) (Fig. 6, C-D), suggest that ZmGT-3b may function upstream of ZmHY5 or as an interacting light- and growth-related partner for ZmHY5, to coordinately regulate the transcription of photosynthesis-related genes during young seedling growth. The interaction of ZmGT-3b with other TFs might be important for its diverse regulatory functions.

### *ZmGT-3b* Knockdown Induces Defense Responses without Immune Activation by Regulating Lignin biosynthesis and Defense-related Gene Expression

Diminished growth is thought to be an integral facet of induced resistance and a molecular mechanism involved in the crosstalk between growth and defense responses. This process involves optimization of the temporal and spatial expression of defense genes. Pathogen infection usually affects primary metabolism, reduces plant growth, limits photosynthesis, and modifies secondary metabolism towards defense responses (Guo et al., 2018). Consistently, *ZmGT-3b* knockdown led to diminished seedling growth and reduced Pn, but improved disease resistance and drought tolerance, and increased lignin contents in young seedlings (Fig. 2, 3, 8). Moreover, the upregulated DEGs induced by *ZmGT-3b* knockdown were enriched in GO terms involved in the biosynthesis of secondary metabolites (especially biosynthesis of phenylpropanoid), including *PR*s and lignin biosynthesis-related genes, without immune activation; *F. graminearum* inoculation induced similar transcriptome reprogramming in young seedlings (CKi/CK) (Fig. 5, C-D, Supplemental Fig. S4).

Many upregulated genes encoded multiple members of the MYB, bZIP, WRKY, NAC, ERF, and bHLHs TF families (Supplemental Fig. S6). Some members of these TF families are well-known regulators of lignin biosynthesis, such as NAC-MYB-GRN, which regulate lignin biosynthesis in both dicot and monocot species (Yoon et al., 2015). Five maize MYB TFs (ZmMYB2, ZmMYB8, ZmMYB31, ZmMYB39, ZmMYB42) function in lignin biosynthesis by controlling *ZmCOMT* expression (Fornale et al., 2006). Overexpression of *ZmMYB167* increased lignin content to 13% in maize without affecting plant growth or development (Bhatia et al., 2019). Besides MYB TFs, some NAC (Zhong et al., 2007) and WRKY TFs also regulate lignin biosynthesis by modulating the expression of cell wall synthesis-related genes (Gallego-Giraldo et al., 2016).

The plant cell wall is a highly organized structure composed of lignin, cellulose, semi-cellulose, pectin, proteins, and aromatic substances. Cell wall biosynthesis requires the coordinated action of numerous enzymes that are often coordinately regulated both spatially and temporally by specific TFs (Liu et al., 2018; Ohtani and Demura 2019).

Increasing lignin contents via activation of the immune response is a conserved basal defense mechanism in plants, allowing defense-induced lignification to be used as a biochemical marker of an activated immune response (Dixon and Barros 2019; Vanholme et al., 2019). In addition to well-known enzymes involved in monolignol biosynthesis, DPs, PODs, and LACs are components of the lignin polymerization machinery (Liu et al., 2018). Many genes encoding these enzymes are induced by various biotic and abiotic stresses, highlighting their roles in the biosynthesis of defensive lignin and/or strengthening of the cell wall via lignin deposition in response to stress (Paniagua et al., 2017).

*ZmGT-3b* knockdown led to the significant upregulation of a subset of secondary metabolite biosynthesis-related genes, especially genes encoding lignin biosynthesis enzymes, including PALs, 4CLs, HCTs, CoCOMTs, CCR, COMTs, LACs, DPs, PODs, and CASPs, in *GT-KD* versus CK seedlings (Fig. 8, Supplemental Fig. S6). The mutation of *ZmCOMT*, encoding a caffeic acid O-methyl transferase, leads to the brown *midrib3* phenotype (Fornale et al. 2006). *ZmCOMT* transcript levels were high in maize seedlings, with FPKM values over 1,200; its transcript level increased 1.57-fold in *GT-KD* versus CK seedlings. *ZmMYB19/Zm1* is an ortholog of the *Arabidopsis* SG3-type R2R3-MYB genes *MYB58* and *MYB63*. These TFs transactivate the promoters of monolignol pathway genes, and their overexpression specifically activates the monolignol pathway and lignin accumulation at the expense of biomass production (Zhou et al., 2009). Among the 18 MYB genes with significantly upregulated expression in *GT-KD* versus CK seedlings, *ZmMYB19/Zm1* expression increased 2.04-fold in response to *ZmGT-3b* knockdown (Supplemental Fig. S6). The upregulation of these genes might contribute to the increased lignin and ASL contents of *GT-KD* seedlings (Fig. 8A).

Lignin biosynthesis is coordinately regulated with the biosynthesis of other cell wall components. The deposition of cellulose in cell walls is vital for controlling cell growth (Yoon et al., 2015; Liu et al., 2018; Ohtani and Demura 2019). Consistent with the retarded growth of young *GT-KD* seedlings, the cellulose and semi-cellulose contents were also higher in these seedlings compared with CK seedlings. Indeed, the expression levels of all 22 Cesa-encoding genes were higher *GT-KD* versus CK seedlings, although not all differences were significant (Fig. 8E). These results suggest that *ZmGT-3b* knockdown induces defense responses without activating the immune system by transcriptionally regulating basal defense- and lignin biosynthesis-related genes in maize seedlings. Therefore, ZmGT-3b might be a negative regulator of biological processes related to plant defense responses, thus acting as a molecular hub connecting developmental/environmental signals with lignin biosynthesis during maize seedling growth.

### *ZmGT-3b* Plays a Role in Drought Tolerance

Drought severely limits crop productivity worldwide. Drought causes plant dehydration by disrupting cellular osmotic equilibrium, ultimately leading to various physiological and metabolic disorders such as damaged photosynthetic activity and excess ROS production. Plants have evolved various mechanisms to protect themselves from drought stress or to tolerate dehydration. Such mechanisms include the biosynthesis of various low molecular weight osmotic-protective compounds, maintaining cell water content with ion accumulation (Mak et al., 2014), reinforcing the cell wall (Tenhaken,2015), and detoxifying ROS (Zhao et al. 2018). The biosynthesis and accumulation of osmotic-protective compounds is an energy- and assimilate-consuming process. Increasing cytosolic ion concentrations by taking up inorganic ions from the environment via transporters and channels is a much more cost-effective method for intracellular osmotic adjustment in plants (Conde et al. 2011).

Some ion transporters are involved in regulation of ion homeostasis and balancing ROS production. Potassium (K^+^) is the most abundant cation and an essential element for plants. K^+^ is critical for the adaptive responses of plants to various abiotic or biotic stresses, including drought stress, as increased K^+^ uptake confers higher levels of drought tolerance (Feng et al.,2016; Cai et al.,2019). Phosphorus (P) is an indispensable nutrient for plant growth and development, as it is a constituent of many important molecules such as nucleic acids, phospholipids, and ATP (Guo et al., 2015). Cu is critical for electron transport and for scavenging ROS produced in chloroplasts during photosynthesis under stress conditions (Boutigny et al., 2014).

*ZmGT-3b* knockdown significantly increased K, P, and Cu contents in *GT-KD* seedlings, and the corresponding transporter genes were upregulated in these seedlings compared with CK seedlings (Supplemental Fig. S7). Two sulfate transporter genes were also significantly upregulated in *GT-KD* seedlings. Sulfate transporters are important for plant drought and salinity tolerance, as sulfate accumulation in leaves enhances ABA biosynthesis, leading to stomatal closure (Gallardo et al., 2014). The increased accumulation of sulfate, P, K, and Cu in the absence of stress might contribute to the enhanced drought tolerance of *GT-KD* seedlings. Fe is involved in various chelation and oxidation/reduction steps that affect ROS production because it is a component of all photosystems and a critical redox-active metal ion in photosynthetic electron flow. Therefore, Fe homeostasis must be fine-tuned in plants (Briat et al., 2007). Al is toxic to plants and seriously affects plant growth and productivity (Yin et al. 2010). Finally, since lignin is a component of the cell wall and the first barrier for metal ions, lignin biosynthesis is associated with heavy metal absorption, transport, and tolerance in plants. Lignin binds to multiple heavy metal ions and reduce their entry into the cytoplasm due to its numerous functional groups (e.g., hydroxyl, carboxyl, and methoxyl) (Dalcorso et al., 2010). The significantly reduced Al and Fe contents in *GT-KD* versus CK seedlings might be associated with increased lignin biosynthesis in response to *ZmGT-3b* knockdown, although one vacuolar iron transporter gene was significantly upregulated in these seedlings, and no Al-related transporter genes were identified in *GT-KD* seedlings.

Members of the AP2/ERF, MYB, bZIP, and NAC TF families are involved in the transcriptional regulation of genes required for drought tolerance. MYBs are important regulators of both secondary cell wall biosynthesis and abiotic stress tolerance, perhaps linking the abiotic stress response and lignin biosynthesis pathways. Drought induces the expression of many *MYB* genes: 65% of *MYB* genes in rice that are expressed in seedlings are differentially regulated under drought stress (Katiyar et al., 2012), and different MYB TFs are involved in one or more drought response mechanisms (Baldoni et al. 2015). *Betula platyphylla* plants overexpressing *BplMYB46* showed increased salt and osmotic stress tolerance and enhanced lignin deposition (Guo et al., 2017). Lignin is important for plant survival under abiotic stress, as defense-induced lignification is a conserved basal defense mechanism, and lignin biosynthetic genes are upregulated in various plants under abiotic stress. Lignin helps maintain osmotic balance in the cell and protects membrane integrity by reducing the penetration of water into the plant cell and hampering transpiration, and lignin biosynthesis increases under drought stress (Mourasobczak et al., 2011). MYB15 plays a role in the complex regulatory relationship between lignin, growth, and defense (Chezem et al., 2017).

Consistent with our observation of a reduced TR and increased lignin contents in *GT-KD* seedlings (Fig. 4E, Fig. 8A), *ZmGT-3b* knockdown induced the significant upregulation of multiple TF genes in *GT-KD* seedlings, including multiple members of the bZIP, MYB, WRKY, NAC, ERF, and bHLH families (Fig. 7). Considering that these TFs are critical in the complex regulatory relationship between growth, lignin, and defense, their elevated expression might be associated with the improved disease resistance and drought tolerance of *GT-KD* seedlings. Collectively, the reinforced cell wall with increased lignin content, the increased accumulation of various mineral elements, and the significantly upregulated expression of various defense-related TFs might all contribute to the drought tolerance of *GT-KD* seedlings.

### ZmGT-3b Mediates the Growth–defense Tradeoff in Maize Seedlings

Based on our results, we propose a model for the mode of action of ZmGT-3b, as shown in Fig 9. According to this model, under normal growth conditions, light induces the expression of *ZmGT-3b*. ZmGT-3b may act upstream or as an activator of ZmHY5 to activate various photosynthesis-related genes, thereby promoting photosynthesis and seedling growth; ZmGT-3b synchronically acts as a transcriptional repressor of the expression of multiple TF genes, including *MYB*s, *bZIP*s, *NAC*s, *bHLH*s, and *ERF*s, which in turn repress the expression of various defense-related genes. When plants are exposed to pathogen attack, *ZmGT-3b* and *ZmHY5* expression were dramatically decreased, thus relieving the repressive effects of these TF genes and enhancing the expression of defense-related genes, including genes encoding PR-proteins and various enzymes involved in the biosynthesis of secondary metabolites especially lignin, thereby activating the defense response. Reduced *ZmGT-3b* expression also leads to a decrease in photosynthesis-related activities to benefit defense-related biological processes. Therefore, perhaps ZmGT-3b and ZmHY5 coordinately regulate the light response and photosynthesis during maize seedling growth, and the interaction of ZmGT-3b with other TFs might be important in its diverse regulatory functions. In conclusion, we propose that ZmGT-3b functions as a transcriptional regulator that calibrates the balance between plant growth and defense responses by coordinating metabolism during growth-to-defense transitions by optimizing the temporal and spatial expression of photosynthesis- and defense-related genes. Therefore, ZmGT-3b might serve as a molecular hub connecting developmental or environmental signals with secondary metabolite biosynthesis.

**Figure 9.**
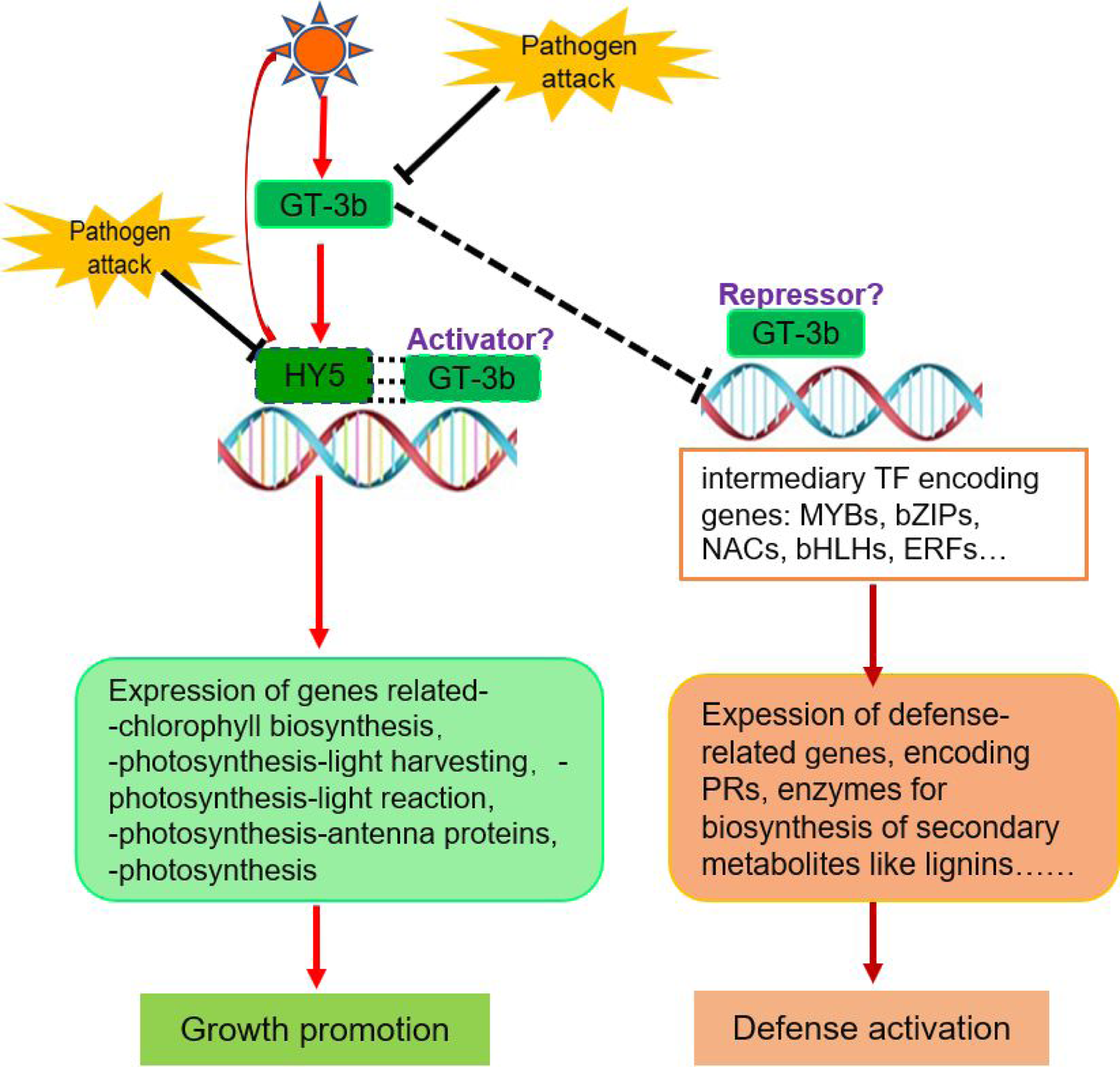
Schematic representation of the mode of action of ZmGT-3b in mediating the growth–defense tradeoff in maize seedlings. Under normal growth conditions, light induces *ZmGT-3b* expression, and ZmGT-3b transcriptionally activates various photosynthesis-related genes, especially *ZmHY5*, to promote photosynthesis and seedling growth; ZmGT-3b synchronically suppresses the expression of various defense-related genes by transcriptionally repressing multiple transcription factor (TF) genes, including *MYB*s, *bZIP*s, *NAC*s, *bHLH*s, and *ERF*s. However, *ZmGT-3b* and *ZmHY5* expression dramatically decreases upon pathogen attack. Thus, photosynthetic activity is limited and the repressive effect on TF genes is relieved, thereby inducing the expression of defense-related genes to activate the defense response. Thick lines indicate direct inhibition by pathogens, while broken lines indicate proposed repression. The dotted lines between ZmGT-3b and ZmHY5 suggest their possible collaboration in regulating the light response and photosynthesis during seedling growth.

## MATERIALS AND METHODS

### Plant Growth, *F. graminearum* Inoculation and Disease Severity Scoring

The fungal pathogen *Fusarium graminearum* preparation and inoculation with *F. graminearum* in the field were done according to Yang et al. (2010); young seedling inoculation on primary roots was done according to Ye et al. (2013); and disease severity scoring was done according to Ye et al. (2018). Three replicates were set for each genotype with about 25 plants per replicate. The primary roots with typical symptoms were scored at 48 hours after inoculation (hai). The length of the 7-days after germination (DAG) young seedling primary roots cultured with paper-rolling were measured and used for comparing seedling root growth rate, and shoot growth rate was measured with the soil-growth young seedlings at 12 or 15 days after germination. The seedlings (CK and *GT-KD*) were cultured in controlled growth room conditions of 28/22°C (day/night) at a light intensity of 500 μmolm^−2^ s^−1^ (16-h-light/8-h-night) and 40–50% relative humidity. Statistical analysis was conducted using Student’s t-test between the control (CK) and mutant *GT-KD* seedlings to determine statistical significance in three independent experiments.

### Generation of the Transgenic Knockdown Lines of *ZmGT-3b*

According to the maize genome sequence RefGen V3.22 in 2014, the third exon (encoding 149aa) of *ZmGT-3b* was obtained by RT-PCR and was put under the control of maize *Ubiquitin* promoter in a *pBXCUN*-derived binary vector to generate *pUbi::cZmGT-3b* (for primer sequences see Supplemental Table S2), the construct was transformed into *Agrobacterium* strain *EHA105*, and then into the immature embryos of the *Zea mays L*. B73-329 inbred lines (used as the control, CK, in the afterward experiments). Six transgenic events of the construct were obtained, self-crossed and enough homogenous seeds were harvested.

### Plasmid Construction and Subcellular Localization Analysis

The full coding sequence (cds) of *ZmGT-3b* was obtained by RT-PCR with gene-specific primers (designed according to the B73 reference genome RefGen V4.32, primer sequences see Supplemental Table S2) amplified with reverse-transcription cDNA templates from maize young seedlings at 7-DAG. Subsequently, the sequenced clone was used for constructing *p1300-35S*: *ZmGT-3b-GFP* vector without the stop codon, with a *pCAMBIA1300*-derived binary vector by the introduction of the cds to fuse to GFP driven by the Camv35S promoter. *Agrobacterium* strain *EHA105* containing *p1300-35S*: *ZmGT-3b-GFP* vector was cultured at 28°C overnight. Bacteria cells were harvested by centrifugation and resuspended with buffer (10mM MES, pH 5.7, 10mM MgCl2, and 200 mM acetosyringone) at OD_600_ = 0.6. Leaves of 3-week-old soil-grown *N. benthamiana* were infiltrated with *Agrobacterium* cultures carrying a binary vector *p1300-35S*:*:GFP* or *p1300-35S*::*ZmGT-3b-GFP* expressing GFP or ZmGT-3b-GFP. The plants were incubated under 16h light/8h dark at 25°C in growth chambers. The GFP fluorescence signals were detected 2-days post-infiltration (dpi). The *Agrobacterium* cultures carrying the binary vectors were also used to transform onion epidermal cells. Fluorescence images were examined and taken with LSM 880 confocal laser microscope systems and images were processed using LSM microscope imaging software. The excitation laser of 488 nm was used for imaging GFP signals.

### Maize Seedling Drought Stress Analysis

After germination, the seedlings (CK and *GT-KD*) were cultured in controlled growth room conditions of 28/22°C (day/night) at a light intensity of 500 μmolm^−2^ s^−1^ (16-h-light/8-h-night) and 40–50% relative humidity. They were grown under well-watered conditions by maintaining soil water content close to field capacity for approximately 10 days until drought treatment. Drought stress (cessation of watering) was imposed on the growing seedlings after two leaves; that is, at about 10 days after sowing by withdrawing water supply and keeping the plants under observation for the following 15 days, when indications of severe withering symptom (all the leaves turned soft, drooping and dried) were visible in almost all of the CK seedlings, then the seedlings were re-watered for 6 days.

Measurements were made at day 15 after the stress treatment and at day 6 following the start of re-watering. For each treatment, the normally irrigated plants were used as controls. The phenotypes and physiological indexes of the seedlings were detected, and the number of surviving seedlings and the total seedling number to obtain the survival rate were checked. The water loss rate test was done with the third leaf of well-grown seedlings at 15 DAG; the leaves were collected and put on to flat pallets, separately and individually under the same environment as the seedlings were growing. The weight of the five leaves was measured as a group at given time; the weight of the lost water was obtained by subtracting this weight from the fresh weight, and the water loss rate was calculated as a percentage of the weight of the lost water to the initial fresh weight of the given group. Data are means ± SD of three replicates.

### Analysis of Cell Wall Components, Cellulose and Lignin Content

The 12-day-old seedlings were harvested, dried and homogenized to a fine powder using a mixer mill (MM400, Retsch Technology, Haan, Germany) at 25 Hz for two minutes. One-hundred milligrams of powdered seedling tissue was sequentially ultrasonicated for 15 min in a mixture of twice with 1 mL methanol, twice with phosphate buffered saline pH (7.0) containing 0.1% (v/v) Tween 20, twice with 1 mL 95% ethanol, twice with 1 mL (1:1) chloroform: methanol and twice with 1 mL acetone. The samples were centrifuged at 16,000 g for 10 min and the pellets dried at 50°C. The remaining cell wall extract was used for determination of total lignin content. The lignin content of seedlings was quantified using the acetyl bromide soluble lignin method. Seedling tissue was macerated in 72% (v/v) sulfuric acid for 2 h, diluted with 112 mL deionized water, and thereafter autoclaved at 121°C for 1 h. The acid-insoluble lignin was quantified using pre-weighed medium coarse-ness glass crucibles, while a UV/VIS Spectrometer was used to determine the acid-soluble lignin content at 205 nm with an ultraviolet spectrophotometer (TU-1901). To hydrolyze the cell wall polysaccharides, 10 mg of destarched sample was mixed with 200 µl 72% sulfuric acid and incubated at 60°C for 1h, and then the sulfuric acid was diluted to 3% with distilled water for hydrolysis at 121°C for 1 h. After cooling to room temperature, the supernatant was collected, erythritol was added as an external standard, and then was neutralized with barium carbonate. The sugars in the supernatant were separated using an SP0810 column (Shodex) on a UHPLC system (Agilent-1260). The content of the detected sugars was calculated based on standard curves of glucose, xylose, mannose, galactose and arabinose. Error bars indicate SD of three biological replicates.

### Leaf Chlorophyll Content, Net Photosynthetic Rate and Transpiration Rate Analysis

Leaf chlorophyll content was measured from the latest expanded leaves of the CK and *GT-KD* seedlings grew at 12- or 15-day after germination (DAG), with a SPAD meter (SPAD-502 Plus, Konica Minolta, Inc. Tokyo, Japan) under a saturating actinic light (660 nm) with an intensity of 1100 µmol m^-2^s^-1^. The middle widest part of the latest expanded leaf of every seedling, that is the second leaf of the12-DAG seedlings and the third leaf of the 15-DAG seedlings, was used for SPAD value measurement. The net photosynthetic rate (Pn) and transpiration rate were measured from the latest expanded leaf (the third leaf) of the 15-DAG seedlings (with three leaves and a heart leaf) with a portable LI-6400XT Portable Photosynthesis System (LI-COR, USA), recorded at a saturating actinic light (660 nm) with an intensity of 1100 µmol m^-2^s^-1^, at the time from 09:00 to 12:00 in the morning. All measurements were conducted on the middle part of the latest expanded leaves following the manufacturer’s instructions. Five replicates were randomly taken for each genotype.

### RNA Extraction and Transcriptome Sequencing

To compare the transcriptomes between *GT-KD* and CK (Control, B73-329) with or without inoculation, the inoculated (the seedlings were inoculated at 5-DAG and sampled at 18 h after inoculation) and the non-inoculated *GT-KD* and CK seedling samples (whole seedlings without the kernel) were collected at the same time (at 18 hai) and used for RNA extraction and deep sequencing. Total RNA was extracted using RNAiso Plus (Takara Bio) according to the user manual. Three micrograms of total RNA from each sample were used for transcriptome sequencing at Novogene (http://www.novogene.com/). Sequencing was performed on each library to generate 100-bp paired-end reads employing the high throughput sequencing platform highseq3000. Read quality was checked using FastQC and low quality reads were trimmed using Trimmomatic version 0.32 (http://www.usadellab.org/cms/?page=trimmomatic). The clean data for each sample amounted to ∼6 Gb. The clean reads were aligned to the masked maize genome database for mapping, calculation, and normalization of gene expression (the updated *Z. mays* B73 reference genome AGPv4, http://ensembl.gramene.org/Zea_mays/Info/Index).

Calculation and normalization of gene expression were based on the reads per kilobase per million mapped reads calculation. Differentially expressed genes (DEGs) were defined with a *P*-value of 0.05, with statistical correction using Benjamini–Hochberg false discovery rate (FDR) of 0.05 in cuffdiff. Parameters used for screening the DEGs were the fold change (FC) of the expression level (FC ≥ 2 or FC ≤ 0.5 under *P*-value ≤ 0.05 and FDR ≤ 0.05), compared with the expression level in the control transcriptome. Defining DEG and cluster analysis was done with the software Cluster 3.0. For GO term analysis, significantly differentially expressed genes were analyzed using AGRIGO (http://bioinfo.cau.edu.cn/agriGO/ between the tested conditions (Trapnell et al., 2010).

### Quantitative real time Reverse Transcription PCR (qRT-PCR) Analysis

Real-time RT-qPCR was used to check the relative expression level and validate the RNA-seq results for those light- and defense-related genes that showed different fold changes in expression level between the *GT-KD* and CK seedlings. Primers were designed with Primer Express Software, according to standard parameters for real-time RT-qPCR assays (Bio-Rad Laboratories, Hercules, CA, U.S.A.). For qRT-PCR experiments, total RNA was extracted from young seedling tissues (the CK and different *GT-KD* lines from three different transgenic event were collected and used for total RNA extraction) at 6-DAG with RNAiso Plus (Takara Bio) according to the user manual. First-strand cDNA was synthesized using about 1µg of total RNA and RT Master Mix with gDNA Remover (Takara Bio), which contains M-MLV reverse transcriptase, oligo(dT) primer and random hexamer primer. qRT-PCR analyses were conducted using PowerUp^TM^ SYBR Green Master Mix (Applied Biosystems, Carlsbad, CA, USA) on an ABI 7500 thermocycler (Applied Biosystems). The qRT–PCR amplification reactions were performed in a total volume of 20 µl for each reaction, which contained 10 µl SYBR Green Realtime PCR Master Mix, 1µl forward and reverse primers (10 µM), 1µl cDNA dilution products and 8 µl H_2_O. The maize *GAPDH* (accession no. X07156) gene was employed as the endogenous control. The sequences of all the primers for qRT–PCR analysis are given in Supplemental Table S2. PCR was performed with the following conditions: 94℃ for 2 min; 40 cycles of 94℃ for 30 s, 60℃ for 30 s, 72℃ for 30 s. The relative expression levels were calculated using the relative quantization according to the quantification method (2^-ΔΔCt^) (Livak and Schmittgen, 2001) and plotted with standard errors. The variation in expression was estimated using three biological replicates independently by comparative reverse transcription-quantitative PCR (RT-qPCR).

## Statistical Analyses

Statistical analysis was performed using the paired Student’s t-test. All values represent the mean ± SD. Asterisks indicate significant differences (\**P* < 0.05, \*\**P* < 0.01, \*\*\**P* < 0.001). All experiments were independently repeated at least three times.

## Supplemental Figure Legends

**Supplemental Figure S1. Expression profile of ZmGT-3b.** A, *ZmGT-3b* expression was rapidly and dramatically decreased after inoculation in both the resistant *qRfg1* allele (R-NIL) or the susceptible *qRfg1* allele (S-NIL), which were developed from a QTL *qRfg1* on chromosome 10 that could explain 36.6% of the total variations of maize resistance to *Fusarium graminearum* induced stalk rot (Yang et al. 2010). RCK (SCK) is the primary roots of R-NIL (S-NIL) without inoculation, R6, R18, R48 (S6, S18, S48) was the primary roots of R-NIL (S-NIL) inoculated after 6h,18h and 48h, respectively (Ye et al 2013). B and C, *ZmGT-3b* expressed highly in normal seedling roots, and only expressed in a few kinds of young tissues, such as primary roots, ear primordium (2-8µm), embryo at 20 DAP and presheath et al (Chen et al, 2014; Johnston 2014; Maize atlas Stelpflug 2015; Walley 2016).

**Supplemental Figure S2. Maize mature kernel weight and plant height analysis.** CK is the wild type plants (B73-329), G3, G6 and G7 were different transgenic maize seedlings with *ZmGT-3b* knock-down, all the plants were grew in the field under normal condition in Beijing, 2018. 2w/4w was 2/4 weeks and MP was mature plant. Three independent repeats were done in the field. Values are the mean ± SD. The asterisk * represents a significant difference at *P* < 0.05 (according to a paired Student’s t-test); NS, not significant.

**Supplemental Figure S3. The relative expression fold of the photosynthesis-related genes from transcriptome sequencing with *ZmGT-3b* knock-down (GT) and CK (B73-329) maize seedlings at 7-DAG, with (CKi, GTi) or without inoculation (CK, GT).** *ZmGT-3b* knock-down dramatically down-regulated genes associated with photosynthesis-related functional categories.

**Supplemental Figure S4. Significant commonalities exist in the transcriptional responses between the *ZmGT-3b* knockdown (GT) and inoculated control plants (CKi).** *ZmGT-3b* knockdown (GT) induced similar transcriptional reprogramming to the inoculated control plants (CKi), to enhance defense related functional categories and to up-regulate expression of defense-related genes.

**Supplemental Figure S5. Comparative transcriptome composition between the various transcriptome pairs.** A, Inoculation induced 1340 genes to be differently expressed in *GT-KD* seedlings (GTi/GT), while 1049 genes were differently expressed in the inoculated two genotypes (GTi/CKi). B, 254 differentially expressed genes (DEGs) were shared between GT/CK and GTi/CKi, while 462 DEGs were shared between CKi/CK and GTi/GT. C-D, Significant commonalities exist in the transcriptional responses between the non-inoculated GT/CK and the inoculated GTi/CKi by GO and KEGG analysis. E-F, The transcriptional reprogramming between the inoculated and non-inoculated *GT-KD* seedlings (GTi/GT) was much stronger than that of the inoculated and non-inoculated CK seedlings (CKi/CK).

**Supplemental Figure S6. The relative expression fold of the defense-related genes from transcriptome sequencing with *ZmGT-3b* knock-down (GT) and CK (B73-329) maize seedlings at 7-DAG, with (CKi, GTi) or without inoculation (CK, GT)**. *ZmGT-3b* knockdown dramatically upregulated genes that emphasized in functions in plant defense response to various biotic/abiotic stress, such as POD, PR, CASP and a few TF family member encoding genes, such as bHLH, bZIP, MYB, WRKY, ERF, and NAC,.

**Supplemental Figure S7. Contents of the mineral elements and expression fold of the related transporter encoding genes in maize seedlings.** A, The contents of mineral elements in CK and *GT-KD* maize seedlings at seven day-after-germination (7-DAG). Compared with CK seedlings, the content of Al and Fe was significantly decreased, while the content of Cu, K and P was increased in the *GT-KD* seedlings. DW, Dry weight. Values are the mean ± SD. The asterisk * represents a significant difference at *P* < 0.05 (according to a paired Student’s t-test); NS, not significant. B, The relative expression fold of the related mineral element transporter encoding genes from the transcriptome sequencing with *ZmGT-3b* knock-down (GT) and CK (B73-329) maize seedlings at 7-DAG, with (CKi, GTi) or without inoculation (CK, GT).

**Supplemental Table S1.** The cis-element analysis within 2000bp upstream of *ZmGT-3b* (up), *ZmHY5* (down) starting codon ATG.

**Supplemental Table S2.** All primers used in the experiments.

